# Refinement of *α*-synuclein ensembles against SAXS data: Comparison of force fields and methods

**DOI:** 10.1101/2021.01.15.426794

**Authors:** Mustapha Carab Ahmed, Line K. Skaanning, Alexander Jussupow, Estella A. Newcombe, Birthe B. Kragelund, Carlo Camilloni, Annette E. Langkilde, Kresten Lindorff-Larsen

## Abstract

The inherent flexibility of intrinsically disordered proteins (IDPs) makes it difficult to interpret experimental data using structural models. On the other hand, molecular dynamics simulations of IDPs often suffer from force-field inaccuracies, and long simulations times or enhanced sampling methods are needed to obtain converged ensembles. Here, we apply metainference and Bayesian/Maximum Entropy reweighting approaches to integrate prior knowledge of the system with experimental data, while also dealing with various sources of errors and the inherent conformational heterogeneity of IDPs. We have measured new SAXS data on the protein *α*-synuclein, and integrate this with simulations performed using different force fields. We find that if the force field gives rise to ensembles that are much more compact than what is implied by the SAXS data it is difficult to recover a reasonable ensemble. On the other hand, we show that when the simulated ensemble is reasonable, we can obtain an ensemble that is consistent with the SAXS data, but also with NMR diffusion and paramagnetic relaxation enhancement data.

## Introduction

Intrinsically Disordered Proteins (IDPs) play important roles in a wide range of biological processes including cell signalling and regulation (***Uversky et al., 2005***; ***Das et al., 2015***; ***Snead and Eliezer, 2019***), and their malfunction or aggregation is linked to neurodegenerative diseases such as Alzheimer’s and Parkinson’s diseases. A key, defining property of IDPs is that they do not adopt well-defined, permanent secondary and tertiary structures under native conditions, and their conformational properties are thus best described in statistical terms.

Due to the dynamic nature of IDPs and their inherent conformational heterogeneity, IDPs are not easily amenable to high-resolution characterisation solely through experimental measurements. To characterise their structural and dynamic properties it is often necessary to integrate various biophysical experiments, and particularly nuclear magnetic resonance (NMR) spectroscopy (***Dyson and Wright, 2001***), small angle X-ray scattering (SAXS or SANS) (***Bernado and Svergun, 2012***), circular dichroism (***Chemes et al., 2012***), and single-molecule Förster resonance energy transfer (sm-FRET) (***LeBlanc et al., 2018***) have been widely used to characterise the structural properties of IDPs. For instance, pulsed-field-gradient NMR diffusion and SAXS experiment are especially useful to quantify the level of compaction of the IDP. Techniques such as sm-FRET and NMR paramagnetic relaxation enhancement (PRE) provide distance information between different residues or regions of the IDP (***Dedmon et al., 2005***; ***Eliezer, 2009***). Nevertheless, since most experimental methods only convey ensemble averaged information and are also affected by random and systematic errors, it is difficult to extract directly information on the underlying heterogeneous ensemble of the IDP. To address this problem, theoretical and computational models can be used to extract detailed structural information from these experiments.

Molecular dynamics (MD) simulations that use physics-based force fields may provide high-resolution temporal and spatial information about the structure and dynamics of IDPs. Extensive sampling of a force field with MD simulations can thus be used to generate conformational ensemble of the IDP. The quality of the results, however, depends heavily on the accuracy of the force field employed. For example it has been shown that many earlier generation of force fields produce overly compact conformations for many IDPs (***Piana et al., 2015***). It appears that these force fields fail to accurately describe the solvation of the protein by underestimating protein-water interactions (***Sun and Kollman, 1995***; ***Nerenberg et al., 2012***; ***Best et al., 2014***; ***Piana et al., 2015***). Recently, however, significant advancements have been made to improve force field accuracy and correct the bias towards overly compact conformations (***Best et al., 2014***; ***Piana et al., 2015***; ***Song et al., 2017***; ***Robustelli et al., 2018***). Adding to these issues, the large conformational phase space of IDPs, requires extensive sampling of the protein is in order to generate converged ensembles. To achieve sufficient sampling, and push the sampling capacity of MD simulations, one often employs enhanced sampling methods such as metadynamics (***Barducci et al., 2008***) or parallel-tempering replica exchange (***Sugita and Okamoto, 1999***). Notably, force field and sampling problems are expected to be more severe for longer IDPs.

An approach to address the challenges of force-field accuracy is to combine experimental and theoretical information in order to obtain conformational ensembles of IDPs that agree with experimental measurements. In this way, the simulations are used as a tool to interpret experimental measurements. A number of different approaches have been described and can, roughly, be divided into two different classes in which the experimental data is either (i) used for on-the-fly restraining of a simulation to experimental data, or (ii) post-processing ensembles generated by simulations to match experimental data by reweighting or selection methods. Many different such methods exist and we refer to a recent reviews for additional details (***Cesari et al., 2018***; ***Orioli et al., 2020***).

Because the conformational ensembles are broad and the experimental data often have low information content and may be noisy, in particular Bayesian inference methods (***Box and Tiao, 2011***) and the maximum entropy principle (***Jaynes, 1957***) have emerged as particularly successful frameworks for studying IDPs. In these frameworks, an ensemble generated using a prior model is minimally modified to match the experimentally observed data better. An extension of these frameworks for integrative structural ensemble determination is Metainference Metadynamics (M&M) (***Bonomi et al., 2016a***), that combines multi-replica all-atom molecular dynamics simulations with ensemble averaged experimental data (***Bonomi et al., 2016b***). In the M&M approach, the metainference (***Bonomi et al., 2016a***) part is a Bayesian inference method that allows for the integration of experimental information with prior knowledge of the system from e.g. physics-based force fields, while also dealing with uncertainty and errors as well as conformationally heterogeneous systems. In addition, metainference can be combined with metadynamics (***Laio and Parrinello, 2002***; ***Bonomi et al., 2016b***) to accelerate sampling further. While metainference applies the bias on the fly, other Bayesian formalisms takes as input simulations that were generated without taking the experimental data into account, and subsequently updates this using statistical reweighting. Such approaches include our Bayesian/Maximum Entropy (BME) protocol (***Bottaro et al., 2020***), as well as related methods (***Hummer and Köfinger, 2015***).

Here, we combined ensemble-averaged experimental SAXS data with MD simulations with the aim to achieve structural ensembles of the system which are in agreement with the experimental data. We did so using both metainference and BME. In particular, we used BME to refine ensembles that had previously been generated using MD simulations (***Piana et al., 2015***; ***Robustelli et al., 2018***), while metainference was applied to restrain experimental SAXS data during MD simulations with an implicit solvent model (***Bottaro et al., 2013***). We used the intrinsically disordered protein *a*-synuclein (*αSN*) protein as a model, as this protein has been studied extensively by various experimental methods including SAXS and NMR measurements, and because of the availability of long MD trajectories generated from a range of force fields and water models. *αSN* is a 140 residue long IDP that is primarily expressed in the brain and in its monomeric state is known to be disordered and populate multiple conformational states. *αSN* aggregation into amyloid fibrils is linked to Parkinson’s disease and dementia with Lewy bodies (***Spillantini and Goedert, 2000***; ***Ulusoy and Di Monte, 2013***).

We assessed the quality of existing ensembles before refinement, and the ability of metainference and BME methods to improve them through incorporation of experimental SAXS data, by comparing with independent measurements of the level of compaction (through the hydrodynamic radius, *R_h_*, as probed by NMR) and previously measured paramagnetic relaxation enhancement data (***Dedmon et al., 2005***). We find that the inclusion of SAXS-restraint in the M&M simulation resulted in the generation of a reliable and heterogenous conformational ensemble that also improved the agreement with the NMR diffusion data. The BME reweighting improved the agreement with the experimental data when we applied the approach to simulations with the TIP4P-D water model. For simulations using the TIP3P water model, which were substantially more compact, it was difficult to find a suitably large ensemble compatible with the experimental SAXS data. Together, our result provide insight into how and when experimental SAXS data can be used to refine ensembles of IDPs, and the role played by the force field as a ‘prior’ in these Bayesian/Maximum entropy approaches.

## Methods and Materials

### Experimental data

Human *αSN* for SAXS experiments was expressed, purified and lyophylized as previously described (***van Maarschalkerweerd et al., 2014***). Prior to SAXS data collection, the lyophilized powder was dissolved in PBS (20 mM Na_2_HPO_4_, 150 mM NaCl, pH 7.4) and filtered through a 0.22 *μ*m filter to remove larger aggregates. The final sample concentration before SEC-SAXS was determined by *A*_280_ to be 4.5 mg/mL using an extinction coefficient of 5960 M^−1^ cm^−1^. SAXS data was collected as SEC-SAXS data on beamline P12 (***Blanchet et al., 2015***) operated by EMBL Hamburg at the PETRA III storage ring (DESY, Hamburg, Germany). 50 *μ*L 4.5 mg/mL *αSN* in PBS buffer (20 mM Na_2_HPO_4_, 150 mM NaCl, pH 7.4) was injected on a Superdex 200inc 5/150 GL column with a flowrate of 0.4 mL/min. The column was pre-equilibrated with the running buffer (PBS with 2% (v/v) glycerol). SAXS data were collected at 20 °C, with continuous exposure of 1 s per frame throughout the SEC elution. Data processing was done using CHROMIXS (***Panjkovich and Svergun, 2018***), averaging sample data from the frames in the monomeric peak and subtracting the buffer signal taken from the flow-through prior to the sample elution to obtain the final scattering profile (Fig. S1).

We purified *αSN* for NMR experiments as previously described (***Skaanning et al., 2020***). Translational diffusion constants for *αSN* (50*μ*M) and 1,4-dioxane (0.2% v/v; as internal reference) were determined by fitting peak intensity decay from diffusion ordered spectroscopy experiments (***Wu et al., 1995***), using the Stejskal-Tanner equation as described (***Prestel et al., 2018***). Spectra (a total of 64 scans) were obtained over a gradient strength of 2 to 98%, with a diffusion time (Δ) of 200 ms and gradient length (*δ*) of 3 ms. Diffusion constants were used to estimate the hydrodynamic radius for *αSN* described (***Wilkins et al., 1999***; ***Skaanning et al., 2020***) (Fig. S2).

We used previously measured PRE data obtained by measuring intensity ratios with spin-labels added at five different positions (residue: 24, 42, 62, 87 and 103) (***Dedmon et al., 2005***).

### Bayesian/Maximum Entropy Reweighting of Unbiased MD simulations

We used previously generated ensembles of *αSN* obtained by long timescale MD simulations with different force fields from the CHARMM and Amber families (here abbreviated by C and A, respectively) and water models (***Piana et al., 2015***; ***Robustelli et al., 2018***) (Table 1). The published simulation using Amber ff99SB-*disp* (***Robustelli et al., 2018***) was later found to be affected by interactions with its periodic image, and has here been replaced by a 73 *μ*s long simulation performed using the same setup but in a 160Å box and available directly from D. E. Shaw Research.

**Table 1.** Ensembles analysed and refined. 1 A99SB-uses a modified version of the TIP4P-D water model. 2 CHARMM36 with EEF1-SB was only used for the metainference metadynamics simulations; here ‘force field’ and reweighted’ refers to two different simulations with and without the experimental bias, respectively. 3 Metadynamics simulation time.

| Force field | Water model | Time(μs) | $R_g$ Force field($\text{\AA}$) | $R_g$ Reweighted($\text{\AA}$) | $R_h$ Force field($\text{\AA}$) | $R_h$ Reweighted($\text{\AA}$) |
| --- | --- | --- | --- | --- | --- | --- |
| A12 | TIP3P | 5 | $15.4 \pm 0.1$ | $19 \pm 1$ | $20.8 \pm 0.1$ | $23.0 \pm 0.1$ |
| A99SB-ILDN | TIP3P | 5 | $15.3 \pm 0.2$ | $16.0 \pm 0.3$ | $20.6 \pm 0.3$ | $21.3 \pm 0.3$ |
| C22* | TIP3P | 6 | $17.1 \pm 0.4$ | $23 \pm 1$ | $22.2 \pm 0.3$ | $26.1 \pm 0.5$ |
| A99SB-ILDN | TIP4P-EW | 5 | $17.9 \pm 0.8$ | $24 \pm 1$ | $22.8 \pm 0.6$ | $26.4 \pm 0.6$ |
| C22* | TIP4P-D | 20 | $23.3 \pm 0.6$ | $29.3 \pm 0.9$ | $26.7 \pm 0.3$ | $29.6 \pm 0.4$ |
| A99SB-ILDN | TIP4P-D | 11 | $25.7 \pm 0.1$ | $31 \pm 1$ | $27.2 \pm 0.6$ | $30 \pm 1$ |
| A12 | TIP4P-D | 11 | $29.7 \pm 0.5$ | $34.1 \pm 0.3$ | $29.7 \pm 0.2$ | $32 \pm 0.5$ |
| A03ws | TIP4P/2005 | 20 | $30 \pm 2$ | $34.3 \pm 0.6$ | $29.1 \pm 1.1$ | $32 \pm 1$ |
| A99SB-*disp* | <sup>1</sup> | 73 | $26 \pm 1$ | $31.9 \pm 0.6$ | $27.7 \pm 0.5$ | $30.8 \pm 0.4$ |
| CHARMM36<sup>2</sup> | EEF1-SB | 3.2<sup>3</sup> | $46 \pm 4$ | $35.4 \pm 0.5$ | $38 \pm 3$ | $33.1 \pm 0.5$ |
| **Experiment** | | | $35.5 \pm 0.5$ | | $28.6 \pm 0.7$ | |

We used our Bayesian/Maximum Entropy (BME) protocol (***Bottaro et al., 2020***; ***Ahmed et al., 2020***) to reweight the initial force field ensembles (Table 1) with the experimental SAXS data, thus obtaining ensembles that are in closer agreement to the experimental data. Briefly described, the BME approach is based on a combined Bayesian/Maximum entropy framework, that enables one to refine a simulation using experimental data while also taking into account the potential noise in the data and in the so-called forward model used to calculate observables for the ensemble. The purpose of the reweighting is to derive a new set of weights for each configuration in a previously generated ensemble so that the reweighted ensemble satisfies the following two criteria: (i) it matches the experimental data better than the original ensemble and (ii) it achieves this improved agreement by a minimal perturbation of the original ensemble. When the initial weights in the ensemble are uniform 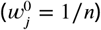, such as when the ensemble has been generated by standard MD simulations, the BME reweighting approach seeks to update the weights, *w_j_*, by minimising the function:

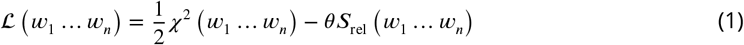

Here, *χ*^2^ quantifies the agreement between the experimental data and the corresponding observable calculated from the reweighted ensemble. 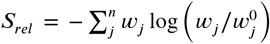 measures the deviation between the original ensemble weights, 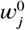, in our case taken as 1/*n*, and the reweighted ensemble weights. Finally, the hyperparameter *θ* tunes the balance between the two terms, and needs to be determined, by evaluating the compromise between the two terms in Equation 1 (***Orioli et al., 2020***). Reweighting and analysis scripts are available at github.com/KULL-Centre/papers/blob/master/2021/aSYN-ahmed-et-al/.

### Metainference Metadynamics

We conducted SAXS-restrained MD simulation using the metainference metadynamics (M&M) method, where we employed the parallel-bias (PBMetaD) flavour of well-tempered metadynamics (***Pfaendtner and Bonomi, 2015***) in combination with the multiple-walkers scheme (***Raiteri et al., 2006***). During the M&M simulation, the SAXS back-calculation step utilises a hybrid-resolution approach, where the SAXS data is calculated on-the-fly using ‘Martini beads’ that are superimposed on the all-atom structures using PLUMED (***Bonomi and Camilloni, 2017***; ***Paissoni et al., 2019***, ****2020****; ***Jussupow et al., 2020***). The approach is particularly efficient as the SAXS back-calculation is calculated using the Debye equation from a coarse-grained model and the excess of electron density in the hydration shell is neglected (***Niebling et al., 2014***; ***Paissoni et al., 2020***). We note here that the Martini model is only used for calculating the SAXS data, and the simulations are performed using an all-atom, implicit solvent model as detailed below.

We used GROMACS 2018.1 (***Abraham et al., 2015***) with PLUMED version 2.4 (***Tribello et al., 2014***) to perform the *M*&*M* simulations. We used the CHARMM36 force field (***Best et al., 2012***) with the EEF1-SB implicit solvent model (***Bottaro et al., 2013***). We used a previously generated structure of *αSN* bound to micelles (***Ulmer et al., 2005***) as starting point for an initial 100-ns long high temperature (500 K) simulation, from which we extracted 64 starting conformations for the multi-replica *M*&*M* simulation. Charged amino acids were neutralised in line with the parameterisation of the EEF1 model (***Lazaridis and Karplus, 1999***; ***Bottaro et al., 2013***), leaving a neutral molecule, and performed a minimisation to a maximum force of 100 kJ/mol/nm. The system was further equilibrated for 20 ns per replica with the metainference bias. For the production simulations the sampling of each replica was enhanced by PBMetaD along with twelve collective variables (CVs) consisting of the radius of gyration and 11 AlphaRMSD CVs to enhance sampling of local backbone conformations (***Tribello et al., 2014***). Gaussians were deposited every 200 steps with a height of 0.1 kJ/mol/ps, and the *σ* values were set to 0.2 *nm* for CVrg and 0.010 for all AlphaRMSD CVs, respectively. We rescaled the height of the Gaussians using the well-tempered scheme with a bias-factor of 20 (***Barducci et al., 2008***).

Because calculation of the SAXS data is limiting in these simulations, we re-binned the experimental SAXS data to a set of 19 SAXS intensities at different scattering vectors, ranging between 0.01 Å^−1^ and 0.20 Å^−1^. Metainference was applied every 10 steps of the simulation. We used

a Gaussian noise model, that applies a single Gaussian per SAXS data-point. The scaling factor between experimental and calculated SAXS intensities was sampled with a flat prior between 0.5 and 2.0 (***Löhr et al., 2017***). We average the estimated metainference weights over a time window of 200 steps; this is done to avoid large fluctuations and prevent numerical instabilities due to too high instantaneous forces (***Löhr et al., 2017***). The Plumed input file is available in the PLUMED-NEST database (***Bonomi et al., 2019***) (plumID:21.003; www.plumed-nest.org/eggs/21/003/).

### Paramagnetic Relaxation Enhancement

Paramagnetic Relaxation Enhancement (PRE) via nitroxide spin-labels has been used extensively to study long-range interactions within IDPs. The measured PRE depends in particular on the distance between a paramagnetic centre and protein nuclei, in this case backbone amides. Because the PRE originates from a dipolar interaction, the observed PRE depends on *r*^−6^, and is thus particularly sensitive to transient, short distances. Because simulations were performed without the spin-labels, and because multiple spin-labels were used to probe the structural ensemble of *αSN*, we used a post-processing approach to estimate the location of the unpaired electron on the nitroxide label. In particular, we used DEER-PREdict (***Tesei et al., 2020***), which is based on a Rotamer Library Approach to place spin labels on the protein, to estimate PRE rates. We calculated and compared results from five paramagnetic labelling positions (residue: 24, 42, 62, 87, 103) in *αSN* (***Dedmon et al., 2005***). Additional details are available in the Supplementary Information and in the DEER-PREdict paper (***Tesei et al., 2020***).

## Results and Discussion

Using *αSN* as an example, we compared conformational ensembles generated either directly using molecular dynamics simulations with a molecular mechanics force field, or the same ensemble refined using SAXS data. We also analysed the results of an approach (M&M) that performs this refinement during the simulation. We thus performed (i) a SAXS-restrained multi-replica simulations using metainference metadynamics and (ii) a reference simulation both using CHARMM36 force field (***Best et al., 2012***) used with the EEF1-SB implicit solvent model (***Bottaro et al., 2013***). Both simulations consisted of 64 replicas, with one simulation using metainference to enforce the agreement with experimental SAXS data, whereas a second, reference simulation did not use experimental restraints and thus sampled the force field only. We also analysed nine previously published multi-*μ*s MD simulations which had been generated using different combinations of proteins force fields and water models (***Piana et al., 2015***; ***Robustelli et al., 2018***) from the AMBER (***Lindorff-Larsen et al., 2010***; ***Hornak et al., 2006***; ***Best and Hummer, 2009***; ***Robustelli et al., 2018***) and CHARMM (***Piana et al., 2011***) families in combination with either standard TIP3P (***Jorgensen, 1981***), TIP4P-EW (***Horn et al., 2004***), TIP4P/2005 (***Abascal and Vega, 2005***) or the TIP4P-D (***Piana et al., 2015***) water model. Table 1 summarises the simulations and below we refer to the prior (not refined) ensemble as the ‘force field’ ensemble and the posterior (refined) ensemble as the reweighted’ ensemble.

### Force Field Accuracy and Sampling

Before the refinement procedure we calculated SAXS intensity curves from each structure in the ensembles using PEPSI-SAXS (***Grudinin et al., 2017***). We also calculated the *R_g_* from the protein coordinates and used them to estimate the hydrodynamic radius (*R_h_*) for each conformation using a previously described empirical relationship (***Nygaard et al., 2017***; ***Ahmed et al., 2020***) (Table 1). The experimental *R_g_* = 35.5 Å was obtained through Guinier analysis of the experimental SAXS curve (see Methods), while the experimental *R_h_* = 29.0 Å was obtained through NMR diffusion measurements (Table 1).

In line with previous observations (***Piana et al., 2015***; ***Robustelli et al., 2018***), the ensembles show very different levels of compaction depending on the force field and, in particular, water model used (Table 1 and Fig. 1). When paired with the TIP3P water model, both the Amber or CHARMM force fields produce very compact conformations and show poor agreement with the experimental value of *R_g_*. On the other hand, when paired with the recently parameterised TIP4P-D water model the force fields give rise to more expanded structures and match the experimental values of *R_g_* and *R_h_* considerably better. The ensemble generated using CHARMM36 with the EEF1-SB implicit solvent model on the other-hand produce more expanded structures (Table 1). Of particular relevance to the reweighting described below it is worth noting how the compact ensembles either do not sample any, or at most very few, structures that are expanded as the *average R_g_* observed in experiment (Fig. 1). This observation already suggests that it will be difficult robustly to derive ensembles that are in agreement with the SAXS data as this in particular is sensitive to the *R_g_*.

**Figure 1.**
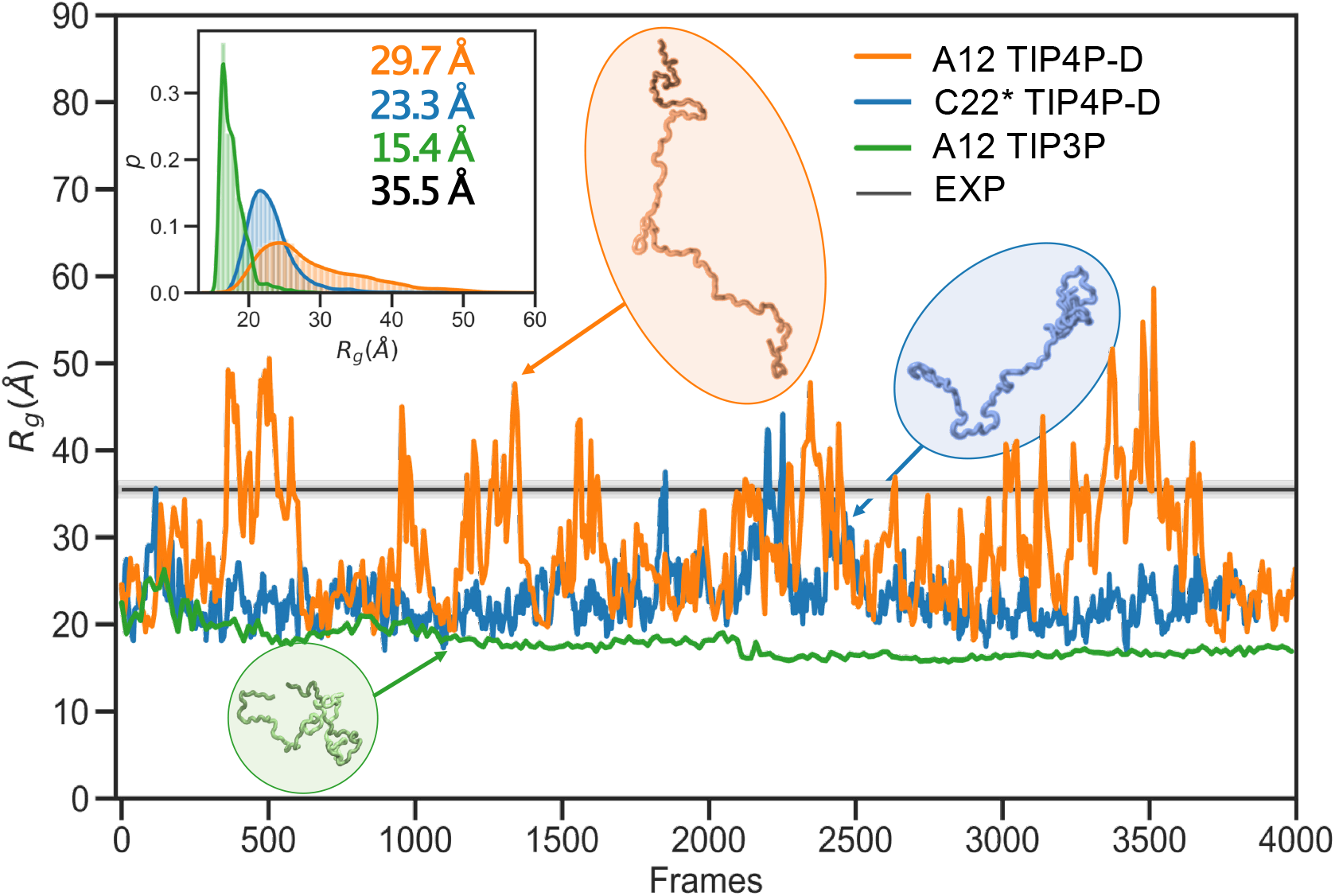
Radius of gyration during simulations with different force fields and water models. As representative examples we show the time-evolution of the radius of gyration for simulations of *αSN* performed with the A12 force field (orange), C22* (blue) and A12 (green) with the TIP4P-D, TIP4P-D and TIP3P water model respectively. The experimental value (black) was obtained from a Guinier analysis of the SAXS data. The orange and blue curves have been smoothed to ease visualization. The insert shows probability densities and averages of *R_g_*. Representative structures with different degree of compaction is also shown. The length of the simulations are 11 *μ*s, 20 *μ*s and 5 *μ*s, respectively, but are shown here on a normalised timescale to make comparisons easier.

### Ensemble refinement using SAXS data

In the following section we exemplify the BME refinement against the SAXS data using two representative combinations of force field and water models, specifically A12 paired with either the TIP3P or the TIP4P-D water model (Figure 2). We also present the results obtained from ‘on-the-fly’ SAXS-restrained simulation with M&M which we compared to an unrestrained simulation with otherwise identical simulation settings (see Methods). Note that while the *R_g_* values for the simulations were calculated using protein coordinates, the experimental value also includes potential contributions from the solvent. The refinement, analysis and plots for the remaining force fields are shown in the supplementary information (Figs. S4–S10).

**Figure 2.**
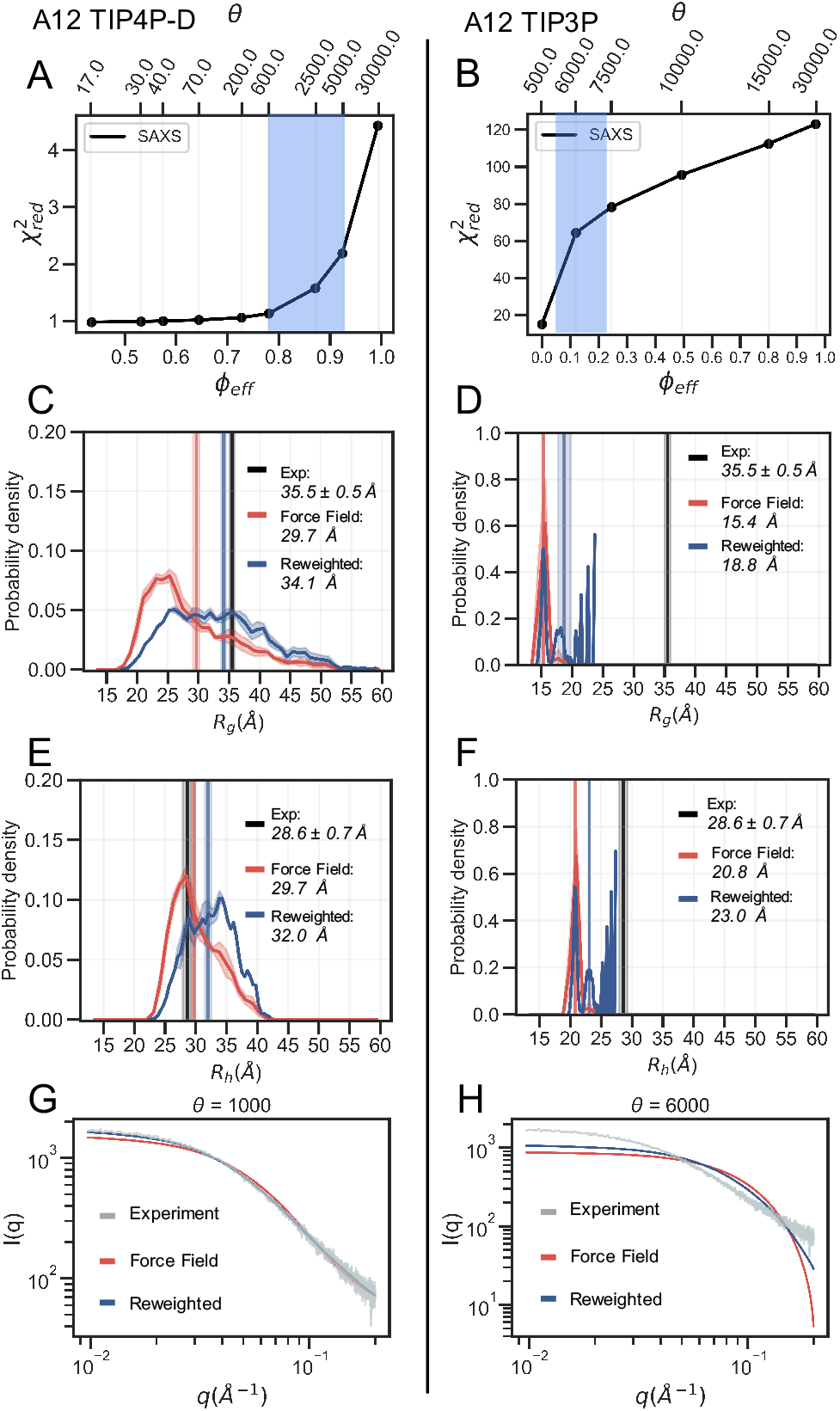
Refinement of two ensembles using BME with SAXS data. SAXS refinement of an ensemble sampled with A12 and either (**left**) the TIP4P-D water model or (**right**) the TIP3P water model. (**A, B**) In the L-curve analysis to select the parameter *θ* we plot *χ*^2^ against *ϕ_eff_*. *θ* balances the prior (force field) and the experimental data, *ϕ_eff_* is the effective number of frames used in the final reweighted ensemble. A value of *θ* is selected from the region marked in blue. We here used *θ* =1000 and *θ* = 6000 for the TIP4P-D ensemble and TIP3P ensemble, respectively. Probability distribution of (**C, D**) *R_g_* and (**E, F**) *R_h_* for the prior (red) and reweighted (blue) ensembles. Solid vertical lines represents the ensemble averaged *R_g_* and *R_h_*. The experimental values are shown in black. The error of the distributions and on the averages (shown as shades) were estimated by block averaging. (**G, H**) Calculated SAXS intensities from the prior ensemble and the reweighted ensembles and are compared to the experimental SAXS data.

The BME procedure works by assigning weights to a previously generated ensemble so as to fit the experimental data better. For BME to successfully reweight an ensemble it is thus required that the initial prior ensemble contains the most relevant conformational states of the protein, such that the ensemble that gives rise to the experimental data is a sub-ensemble of the initial prior ensemble. Consequently, if the sampling is incomplete or the unbiased ensemble is very far away from the true ensemble, it may not be possible to reweight the ensemble to reach a satisfactory agreement with the experiments. An indication that this is occurring is that BME will effectively down-weight most of the structures in the prior ensemble and the posterior ensemble will be dominated by a few structures with large weights. This can in turn be quantified by calculating the (effective) fraction of structures, *ϕ_eff_* = exp(*S_rel_*), that contribute to the ensemble (***Orioli et al., 2020***), so that when *ϕ_eff_* ≈ 1 most of the structures are retained, whereas *ϕ_eff_* ≈ 0 indicates a few structures with very large weights

In the BME reweighting the confidence in the prior ensemble with respect to the experimental data can be tuned by the hyper-parameter *θ* (Eq. 1). One usually does not know the optimal value for *θ* beforehand. Here, we choose *θ* by performing an L-curve analysis (***Hansen and O’Leary, 1993***; ***Orioli et al., 2020***) in which we plot the 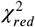 value (quantifying the difference between experiments and calculated value) as a function of *ϕ_eff_*, for different values of *θ* and choose a value corresponding to the ‘elbow’ region (blue region in Fig. 2A and B). The L-curve analysis for the A12 force field paired with TIP4P-D water model, lead us to choose *θ* = 1000, after which the ensemble retains 88% of the initial structures in the final reweighted ensemble, and show much better agreement with the experimental data, indicative by a low 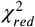 (Fig. 2A). In contrast, the analysis for the TIP3P water model, after reweighting with *θ* = 6000, show that only 12% of the initial structures are used in the final reweighted ensemble in order to achieve significant improved agreement with the experimental data (Fig. 2B). Even at a lower *θ* value there is still a large discrepancy between experimental and calculated SAXS data (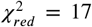 at *θ* = 500). This is a clear example of a poor prior ensemble, which is caused by insufficient overlap between the force field ensemble and that probed by experiment. In fact, the highest value observed (*R_g_* =23 Å) is significantly lower than the experimental value (black). As a consequence, BME ‘throws out’ most of the structures from the initial force field ensemble, and the final reweighted ensemble mainly consist of a few highly weighted structures (Fig. 2D).

The ensemble generated with the TIP4P-D water model (Fig. 2C) contains structures that span a greater range of *R_g_* values, both above and below the experimental value. After refinement the reweighted ensemble is shifted to give greater weight to more expanded structures and bringing the average *R_g_*, substantially closer to the value estimated from the SAXS data. We note here that we do not fit the *R_g_* value but rather the SAXS data. Because the experimental value of *R_g_* (obtained from a Guinier analyses of the data) contains a contribution from the solvent we do not expect a perfect agreement with the average *R_g_* calculated from the protein coordinates (***Henriques et al., 2018***). Indeed, this is one of the reasons why we fit the SAXS data directly rather than the *R_g_*.

The effect of reweighting of the two ensembles can also be seen on the distributions of *R_h_* (Fig. 2E and F). Similarly to *R_g_* distributions, the TIP4P-D ensemble is shifted to give greater weight to more expanded structures (Fig. 2E). As was also evident from the distribution of *R_g_*, the more compact TIP3P ensemble gives rise to a very noisy distribution, because the reweighted ensemble predominantly consist of a few highly weighted structures (Fig. 2F). To illustrate the consequences of reweighting we also compared the calculated SAXS data from the initial force field and reweighted ensembles to the experimental scattering data (Fig. 2G and H). As expected, the refined ensembles show better agreement with experiments, in particular for the A12 paired with TIP4P-D. As agreement between experimental and calculated data is the target for BME this observation again just illustrates that the BME method is indeed optimising agreement.

We repeated these analysis for the remaining combinations of force fields and water models (Figs. S4–S10) and summarise the results by assessing how well the ensembles reproduce *R_g_* and *R_h_* before and after refinement (Fig. 3). We note that the improvement of the *R_g_* observed is due to the use of SAXS data in the refinement, as SAXS intensity curve inherently contains information of the *R_g_*, and that improved agreement with the *R_g_* is thus a sign of the BME approach working rather than a validation of the ensemble.

**Figure 3.**
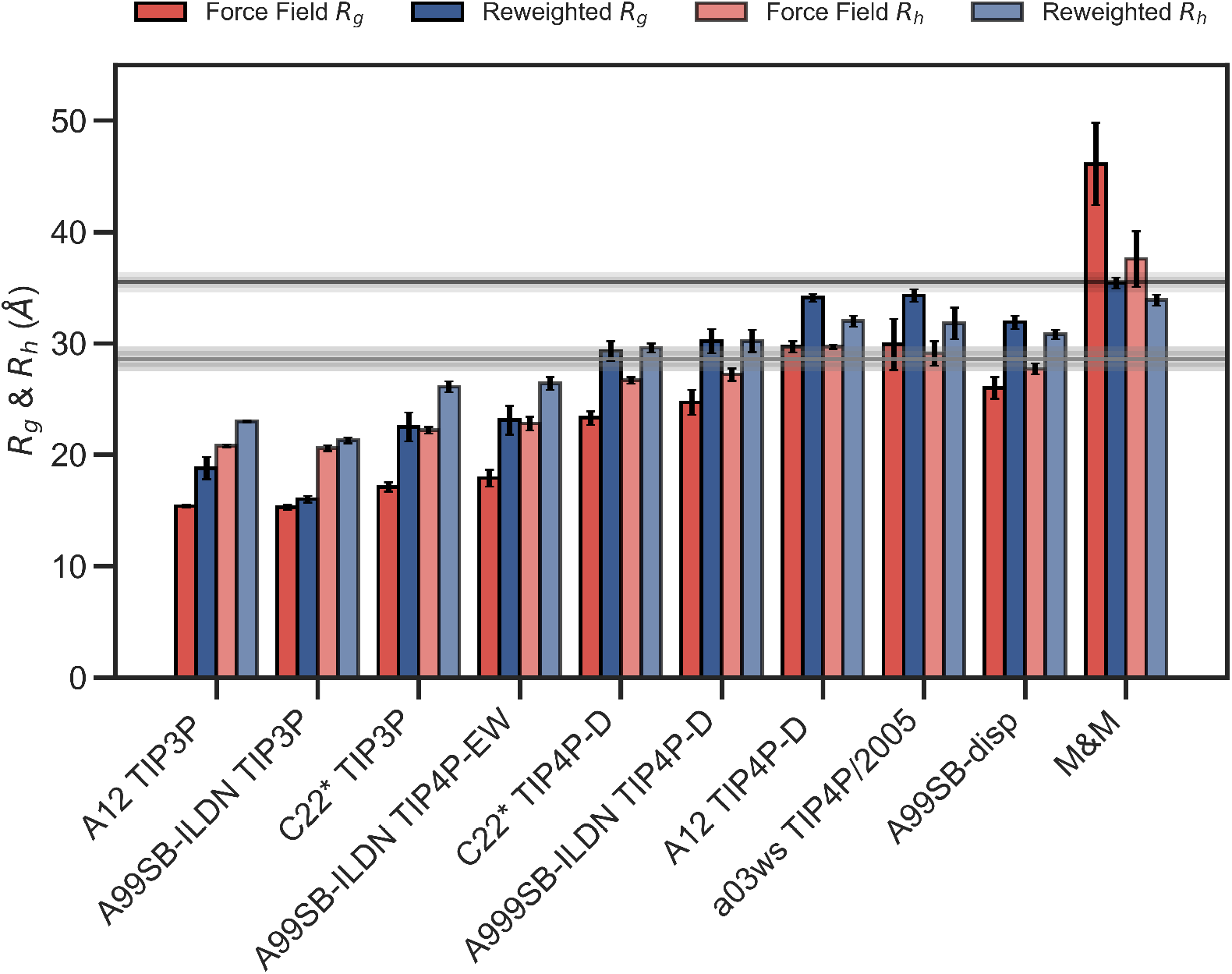
Radius of gyration and hydrodynamic radius calculated from the initial force field ensemble (red) and the experimentally refined ensembles (blue). Experimental values from SAXS (*R_g_* = 35.15Å) and NMR (*R_h_* = 29.0Å) are shown as horizontal lines with the shaded area indicating the error of the experimental values.

To evaluate the effectiveness of the SAXS-restrained M&M simulation we monitored the agreement between the back-calculated and the experimental data over the simulation time by monitoring their correlation rather than the *χ*^2^ (***Paissoni et al., 2020***). Both the SAXS-restrained and the unrestrained reference simulation show a high correlation between back-calculated and experimental data (> 0.98) (Fig. S3A). As expected, the agreement improves substantially when the experimental data is used as a bias in the metainference simulations, confirming the effectiveness of the inclusion of experimental SAXS data (Fig. S3A). Likewise, the average *R_g_*, *R_h_* and the back-calculated SAXS intensity data show improved agreement with the experimental data in the metainference produced ensemble (Fig. 3 and Fig. S3).

In total our analyses show that it is possible to refine MD simulations against SAXS data, though the extent to which agreement can be reached depends on the quality of the input ensemble. For the most compact ensembles we are able to increase the average compaction by fitting to the data, though the average *R_g_* and *R_h_* are still substantially below the experimental values. While the SAXS data (and thus *R_g_*) were used as target values, we also cross-validated with *R_h_* which was not used in the fitting. Here, the picture is less clear. Overall, for the more compact ensembles, fitting the SAXS data lead to improved prediction of *R_h_*. For other ensembles, such as A12 with TIP4P-D, that show good agreement with *R_h_* before reweighting, the agreement became slightly worse after reweighting. Finally, for the most expanded ensemble obtained with CHARMM36/EEF1-SB, agreement with *R_h_* improved after biasing with the SAXS data. We note, however, that the approach we use to estimate *R_h_* from the ensembles is approximate and requires further assessment before these small differences can be interpreted further.

### Validation with PRE data

PRE experiments probe the population-weighted average of the distance (as *r*^−6^) between a paramagnetic centre and protein nuclei, and given the *r*^−6^ dependency is sensitive to the shorter distances even if the populations are small. Here, we compare previously published PREs from spin-labelled *αSN* (***Dedmon et al., 2005***) and back-calculated PRE intensity ratios from five labelling sites, for each of the force field in Table 1, before and after refinement (see also Supporting Information). PRE intensity-ratio profiles from a more expanded ensemble generated using A12 with TIP4P-D (Fig. 4A) and a more compact one generated with A12 with TIP3P (Fig. 4B) show clear differences in agreement before refinement with the SAXS data.

**Figure 4.**
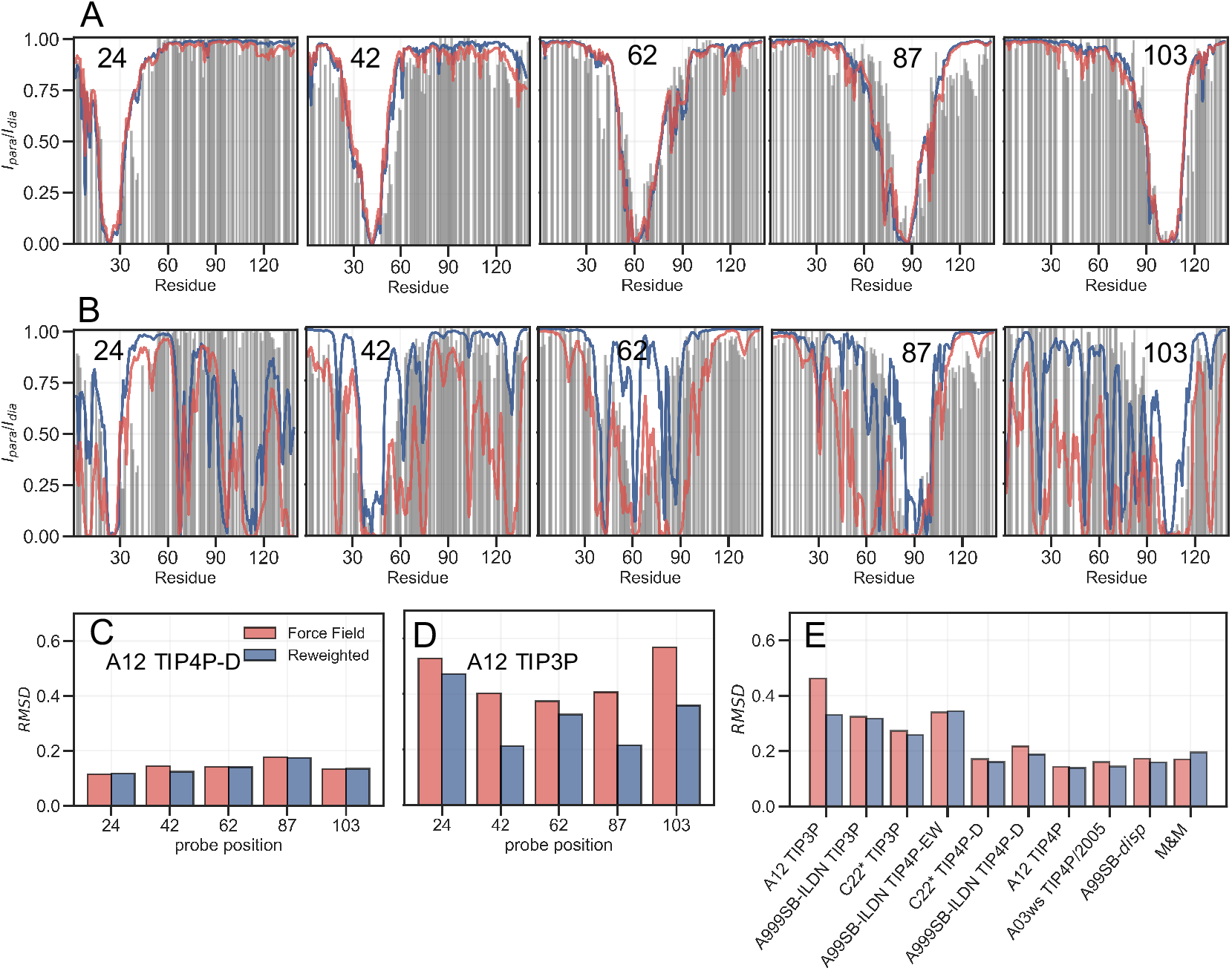
Comparing ensembles to PRE data. We calculated the PRE intensity ratios both from the prior (red) and the reweighted (blue) ensembles and compared to the experimental data (grey). As representative examples we again show results with the A12 protein force field combined with either (A) TIP4P-D or (B) TIP3P water models, and where the location of the spin label probe is denoted in each plot. Experimental intensity ratios slightly exceeding the value 1 were set to 1 in these plots. (C, D) We also calculated the RMSD between the experimental and calculated intensity ratios for each probe and the two force fields both before and after reweighting. (E) Finally, we calculated the RMSD between experiment and calculated values over all probe position for and all force fields in Table 1.

BME refinement leads only to small changes in the calculated PRE data for A12/TIP4P-D, whereas the selection of more expanded structures by applying BME to the ensemble generated with A12/TIP3P leads to more substantial changes as quantified for example by calculating the RMSD between simulation and experimental data (Fig. 4C and 4D). We performed similar calculations and analyses for all ensembles (Figs. S11–S18) and summarize the overall RMSD before and after BME (Fig. 4E). Especially for the force fields paired with TIP3P we observe many of the long-range contacts diminish after reweighting. These results suggest that the reweighting decreases contributions from structures that are too compact, and that the final reweighted ensemble contains more extended structures. In the TIP4P-D ensembles we still observe that some long-range contacts persist even after reweighting and the better agreement is not alone achieved at the cost of a complete elimination long-range contacts; nevertheless, the improvements of the PREs are generally small for these ensembles, and in the case of the metainference ensemble we even observe a small worsening of the agreement.

## Conclusions

We have employed ‘on-the-fly’ or ‘post-facto’ integration between MD simulations and SAXS data *αSN* to derive structural ensembles that are in improved agreement with experiments. These approaches take their outset in a Bayesian framework, and thus the results of the posterior distribution may depend on the choice of the prior. Our results show, in line with previous observations (***Larsen et al., 2020***), clearly that if the prior distribution is a poor model for the experimental data, reweighting becomes noisy. Despite this we find that fitting against SAXS data generally improved or had no effect on the agreement with NMR data (*R_h_* and PREs) that were not target of the optimisation. Thus, the inclusion of a SAXS-restraint in the metainference simulation and the BME refinement showed both methods were able to generate reliable and heterogenous ensemble that maintained good agreement with independent experimental data. We nevertheless also find that the prior used in such protocols are important, and that more robust analyses are obtained with the best priors. Our calculations of *R_h_* and PREs suggest that when the ensembles are ‘far’ away from the experimental data, then improvements driven by the SAXS refinement lead to clear improvements in independent parameters. For ensembles that show better agreement between with the SAXS data to begin with, the picture is less clear. While we on average observe improvements, they are often modest. While some of this is likely because the ensembles are already in reasonably good agreement with experiment, we also suggest that we are observing the limitations of the forward models for calculating SAXS, *R_h_* and PREs. Thus, in addition to improving force fields, future research into finding improved and consistent forward models may be required to provide better models for intrinsically disordered proteins.

## Acknowledgments

We thank A. Kikhney and C. Jeffries for assistance during data collection at the P12 SAXS beamline. We thank D. E. Shaw Research for sharing the molecular dynamics trajectories.

## References

Abascal JL, Vega C. A general purpose model for the condensed phases of water: TIP4P/2005. The Journal of chemical physics. 2005; 123(23):234505.

Abraham MJ, Murtola T, Schulz R, Páll S, Smith JC, Hess B, Lindahl E. GROMACS: High performance molecular simulations through multi-level parallelism from laptops to supercomputers. SoftwareX. 2015; 1:19–25.

Ahmed MC, Crehuet R, Lindorff-Larsen K. Computing, Analyzing, and Comparing the Radius of Gyration and Hydrodynamic Radius in Conformational Ensembles of Intrinsically Disordered Proteins. In: Intrinsically Disordered Proteins Springer; 2020.p. 429–445.

Barducci A, Bussi G, Parrinello M. Well-tempered metadynamics: a smoothly converging and tunable free-energy method. Physical review letters. 2008; 100(2):020603.

Bernado P, Svergun DI. Structural analysis of intrinsically disordered proteins by small-angle X-ray scattering. Molecular biosystems. 2012; 8(1):151–167.

Best RB, Hummer G. Optimized molecular dynamics force fields applied to the helix- coil transition of polypeptides. The journal of physical chemistry B. 2009; 113(26):9004–9015.

Best RB, Zheng W, Mittal J. Balanced protein–water interactions improve properties of disordered proteins and non-specific protein association. Journal of chemical theory and computation. 2014; 10(11):5113–5124.

Best RB, Zhu X, Shim J, Lopes PE, Mittal J, Feig M, MacKerell Jr AD. Optimization of the additive CHARMM all-atom protein force field targeting improved sampling of the backbone *ϕ*, *ψ* and side-chain *χ*^1^ and *χ*^2^ dihedral angles. Journal of chemical theory and computation. 2012; 8(9):3257–3273.

Blanchet CE, Spilotros A, Schwemmer F, Graewert MA, Kikhney A, Jeffries CM, Franke D, Mark D, Zengerle R, Cipriani F, Fiedler S, Roessle M, Svergun DI. Versatile sample environments and automation for biological solution X-ray scattering experiments at the P12 beamline (PETRA III, DESY). Journal of Applied Crystallography. 2015; 48(2):431–443.

Bonomi M, Bussi G, Camilloni C, Tribello GA. Promoting transparency and reproducibility in enhanced molecular simulations. Nature methods. 2019; 16(8):670–673.

Bonomi M, Camilloni C. Integrative structural and dynamical biology with PLUMED-ISDB. Bioinformatics. 2017; 33(24):3999–4000.

Bonomi M, Camilloni C, Cavalli A, Vendruscolo M. Metainference: A Bayesian inference method for heterogeneous systems. Science Advances. 2016; 2(1):e1501177.

Bonomi M, Camilloni C, Vendruscolo M. Metadynamic metainference: enhanced sampling of the metainference ensemble using metadynamics. Scientific reports. 2016; 6:31232.

Bottaro S, Bengtsen T, Lindorff-Larsen K. Integrating molecular simulation and experimental data: a Bayesian/maximum entropy reweighting approach. In: Structural Bioinformatics Springer; 2020.p. 219–240.

Bottaro S, Lindorff-Larsen K, Best RB. Variational optimization of an all-atom implicit solvent force field to match explicit solvent simulation data. Journal of chemical theory and computation. 2013; 9(12):5641–5652.

Box GE, Tiao GC. Bayesian inference in statistical analysis, vol. 40. John Wiley & Sons; 2011.

Cesari A, Reißer S, Bussi G. Using the maximum entropy principle to combine simulations and solution experiments. Computation. 2018; 6(1):15.

Chemes LB, Alonso LG, Noval MG, de Prat-Gay G. Circular dichroism techniques for the analysis of intrinsically disordered proteins and domains. In: Intrinsically disordered protein analysis Springer; 2012.p. 387–404.

Das RK, Ruff KM, Pappu RV. Relating sequence encoded information to form and function of intrinsically disordered proteins. Current opinion in structural biology. 2015; 32:102–112.

Dedmon MM, Lindorff-Larsen K, Christodoulou J, Vendruscolo M, Dobson CM. Mapping long-range interactions in *a*-synuclein using spin-label NMR and ensemble molecular dynamics simulations. Journal of the American Chemical Society. 2005; 127(2):476–477.

Dyson HJ, Wright PE. Nuclear magnetic resonance methods for elucidation of structure and dynamics in disordered states. Methods in enzymology. 2001; 339:258–270.

Eliezer D. Biophysical characterization of intrinsically disordered proteins. Current opinion in structural biology. 2009; 19(1):23–30.

Grudinin S, Garkavenko M, Kazennov A. Pepsi-SAXS: an adaptive method for rapid and accurate computation of small-angle X-ray scattering profiles. Acta Crystallographica Section D: Structural Biology. 2017; 73(5):449–464.

Hansen PC, O’Leary DP. The use of the L-curve in the regularization of discrete ill-posed problems. SIAM Journal on Scientific Computing. 1993; 14(6):1487–1503.

Henriques J, Arleth L, Lindorff-Larsen K, Skepö M. On the calculation of SAXS profiles of folded and intrinsically disordered proteins from computer simulations. Journal of molecular biology. 2018; 430(16):2521–2539.

Horn HW, Swope WC, Pitera JW, Madura JD, Dick TJ, Hura GL, Head-Gordon T. Development of an improved four-site water model for biomolecular simulations: TIP4P-Ew. The Journal of chemical physics. 2004; 120(20):9665–9678.

Hornak V, Abel R, Okur A, Strockbine B, Roitberg A, Simmerling C. Comparison of multiple Amber force fields and development of improved protein backbone parameters. Proteins: Structure, Function, and Bioinformatics. 2006; 65(3):712–725.

Hummer G, Köfinger J. Bayesian ensemble refinement by replica simulations and reweighting. The Journal of chemical physics. 2015; 143(24):12B634_1.

Jaynes ET. Information theory and statistical mechanics. Physical review. 1957; 106(4):620.

Jorgensen WL. Quantum and statistical mechanical studies of liquids. 10. Transferable intermolecular potential functions for water, alcohols, and ethers. Application to liquid water. Journal of the American Chemical Society. 1981; 103(2):335–340.

Jussupow A, Messias AC, Stehle R, Geerlof A, Solbak SM, Paissoni C, Bach A, Sattler M, Camilloni C. The dynamics of linear polyubiquitin. Science advances. 2020; 6(42):eabc3786.

Laio A, Parrinello M. Escaping free-energy minima. Proceedings of the National Academy of Sciences. 2002; 99(20):12562–12566.

Larsen AH, Wang Y, Bottaro S, Grudinin S, Arleth L, Lindorff-Larsen K. Combining molecular dynamics simulations with small-angle X-ray and neutron scattering data to study multi-domain proteins in solution. PLoS computational biology. 2020; 16(4):e1007870.

Lazaridis T, Karplus M. Effective energy function for proteins in solution. Proteins: Structure, Function, and Bioinformatics. 1999; 35(2):133–152.

LeBlanc S, Kulkarni P, Weninger K. Single Molecule FRET: A powerful tool to study intrinsically disordered proteins. Biomolecules. 2018; 8(4):140.

Lindorff-Larsen K, Piana S, Palmo K, Maragakis P, Klepeis JL, Dror RO, Shaw DE. Improved side-chain torsion potentials for the Amber ff99SB protein force field. Proteins: Structure, Function, and Bioinformatics. 2010; 78(8):1950–1958.

Löhr T, Jussupow A, Camilloni C. Metadynamic metainference: Convergence towards force field independent structural ensembles of a disordered peptide. The Journal of chemical physics. 2017; 146(16):165102. doi: 10.1063/1.4981211.

van Maarschalkerweerd A, Vetri V, Langkilde AE, Foderà V, Vestergaard B. Protein/lipid coaggregates are formed during *a*-synuclein-induced disruption of lipid bilayers. Biomacromolecules. 2014; 15(10):3643–3654.

Nerenberg PS, Jo B, So C, Tripathy A, Head-Gordon T. Optimizing solute–water van der waals interactions to reproduce solvation free energies. The Journal of Physical Chemistry B. 2012; 116(15):4524–4534.

Niebling S, Björling A, Westenhoff S. MARTINI bead form factors for the analysis of time-resolved X-ray scattering of proteins. Journal of applied crystallography. 2014; 47(4):1190–1198.

Nygaard M, Kragelund BB, Papaleo E, Lindorff-Larsen K. An efficient method for estimating the hydrodynamic radius of disordered protein conformations. Biophysical journal. 2017; 113(3):550–557.

Orioli S, Larsen AH, Bottaro S, Lindorff-Larsen K. How to learn from inconsistencies: Integrating molecular simulations with experimental data. In: Progress in Molecular Biology and Translational Science, vol. 170 Elsevier; 2020.p. 123–176.

Paissoni C, Jussupow A, Camilloni C. Martini bead form factors for nucleic acids and their application in the refinement of protein–nucleic acid complexes against SAXS data. Journal of Applied Crystallography. 2019; 52(2):394–402.

Paissoni C, Jussupow A, Camilloni C. Determination of Protein Structural Ensembles by Hybrid-Resolution SAXS Restrained Molecular Dynamics. Journal of Chemical Theory and Computation. 2020; 16(4):2825–2834.

Panjkovich A, Svergun DI. CHROMIXS: automatic and interactive analysis of chromatography-coupled small angle X-ray scattering data. Bioinformatics. 2018; 34(11):1944–19464.

Pfaendtner J, Bonomi M. Efficient sampling of high-dimensional free-energy landscapes with parallel bias metadynamics. Journal of chemical theory and computation. 2015; 11(11):5062–5067.

Piana S, Donchev AG, Robustelli P, Shaw DE. Water dispersion interactions strongly influence simulated structural properties of disordered protein states. The journal of physical chemistry B. 2015; 119(16):5113–5123.

Piana S, Lindorff-Larsen K, Shaw DE. How robust are protein folding simulations with respect to force field parameterization? Biophysical journal. 2011; 100(9):L47–L49.

Prestel A, Bugge K, Staby L, Hendus-Altenburger R, Kragelund BB. Characterization of dynamic IDP complexes by NMR spectroscopy. In: Methods in enzymology, vol. 611 Elsevier; 2018.p. 193–226.

Raiteri P, Laio A, Gervasio FL, Micheletti C, Parrinello M. Efficient reconstruction of complex free energy landscapes by multiple walkers metadynamics. The journal of physical chemistry B. 2006; 110(8):3533–3539.

Robustelli P, Piana S, Shaw DE. Developing a molecular dynamics force field for both folded and disordered protein states. Proceedings of the National Academy of Sciences. 2018; 115(21):E4758–E4766.

Skaanning LK, Santoro A, Skamris T, Martinsen JH, D’Ursi AM, Bucciarelli S, Vestergaard B, Bugge K, Langkilde AE, Kragelund BB. The Non-Fibrillating N-Terminal of *α*-Synuclein Binds and Co-Fibrillates with Heparin. Biomolecules. 2020; 10(8):1192.

Snead D, Eliezer D. Intrinsically disordered proteins in synaptic vesicle trafficking and release. Journal of Biological Chemistry. 2019; 294(10):3325–3342.

Song D, Luo R, Chen HF. The IDP-specific force field ff14IDPSFF improves the conformer sampling of intrinsically disordered proteins. Journal of chemical information and modeling. 2017; 57(5):1166–1178.

Spillantini MG, Goedert M. The *α*-synucleinopathies: Parkinson’s disease, dementia with Lewy bodies, and multiple system atrophy. Annals of the New York Academy of Sciences. 2000; 920(1):16–27.

Sugita Y, Okamoto Y. Replica-exchange molecular dynamics method for protein folding. Chemical physics letters. 1999; 314(1-2):141–151.

Sun Y, Kollman PA. Hydrophobic solvation of methane and nonbond parameters of the TIP3P water model. Journal of computational chemistry. 1995; 16(9):1164–1169.

Tesei G, Martins JM, Kunze MB, Wang Y, Crehuet R, Lindorff-Larsen K. DEER-PREdict: software for efficient calculation of Spin-Labeling EPR and NMR data from conformational ensembles. bioRxiv. 2020;.

Tribello GA, Bonomi M, Branduardi D, Camilloni C, Bussi G. PLUMED 2: New feathers for an old bird. Computer Physics Communications. 2014; 185(2):604–613.

Ulmer TS, Bax A, Cole NB, Nussbaum RL. Structure and dynamics of micelle-bound human *α*-synuclein. Journal of Biological Chemistry. 2005; 280(10):9595–9603.

Ulusoy A, Di Monte DA. *α*-Synuclein elevation in human neurodegenerative diseases: Experimental, pathogenetic, and therapeutic implications. Molecular neurobiology. 2013; 47(2):484–494.

Uversky VN, Oldfield CJ, Dunker AK. Showing your ID: intrinsic disorder as an ID for recognition, regulation and cell signaling. Journal of Molecular Recognition: An Interdisciplinary Journal. 2005; 18(5):343–384.

Wilkins DK, Grimshaw SB, Receveur V, Dobson CM, Jones JA, Smith LJ. Hydrodynamic radii of native and denatured proteins measured by pulse field gradient NMR techniques. Biochemistry. 1999; 38(50):16424–16431.

Wu D, Chen A, Johnson CS. An improved diffusion-ordered spectroscopy experiment incorporating bipolargradient pulses. Journal of magnetic resonance, Series A. 1995; 115(2):260–264.

