## Supporting Text and Figures for "Refinement of *α*-synuclein ensembles against SAXS data: Comparison of force fields and methods"

### Supporting Information for: Refinement of $\alpha$-synuclein ensembles against SAXS data: Comparison of force fields and methods

Mustapha Carab Ahmed,[†] Line K. Skaanning,[‡] Alexander Jussupow,[¶] Estella A. Newcombe,[†] Birthe B. Kragelund,[†] Carlo Camilloni,[¶,§] Annette E. Langkilde,[‡] and Kresten Lindorff-Larsen[*,†]

†*Structural Biology and NMR Laboratory & Linderstrøm-Lang Centre for Protein Science, Department of Biology, University of Copenhagen, Denmark*

‡*Department of Drug Design and Pharmacology, University of Copenhagen, Denmark*

¶*Department of Chemistry and Institute for Advanced Study, Technical University of Munich, Germany*

§*Dipartimento di Bioscienze, Università degli studi di Milano, Italy*

### Calculation of Experimental Observables from ensembles

#### Small-angle X-ray scattering (SAXS)

We used Pepsi-SAXS[1] to calculate the SAXS intensity profiles for the different ensembles. As Pepsi-SAXS has free parameters for the solvation layer we used a two-step procedure to determine them at to minimise the risk overfitting by selecting different parameters for each structure.[2,3] First, we fitted the parameters independently to each structure. We then calculate the average for each of the resulting fitted values, and re-ran Pepsi-SAXS with these parameters fixed to those averages.

#### Calculation of Rg and Rh

For each of the ensembles we calculate $R_g$ and $R_h$ as recently described.[4] For a given configuration of a protein, the $R_g$ may be calculated from the protein coordinates as the mass-weighted root mean distance to the centre of mass:

$$R_g = \left( \frac{\sum_i \|\mathbf{r}_i\|^2 m_i}{\sum_i m_i} \right)^{\frac{1}{2}} \tag{1}$$

Where $m_i$ is the mass of atom $i$, $\mathbf{r}_i$ is the position of atom $i$ with respect to the center of mass of the molecule.

We calculated the ensemble averaged radius of gyration ($\langle R_g \rangle$) as:

$$\langle R_g \rangle = \left\langle R_g^2 \right\rangle^{1/2} \tag{2}$$

We estimated $R_h$ through an empirical relationship between $R_g$ and $R_h$[5] :

$$\frac{R_{\mathrm{g}}}{R_{\mathrm{h}}} (N, R_{\mathrm{g}}) = \frac{\alpha_1 \left( R_{\mathrm{g}} - \alpha_2 N^{0.33} \right)}{N^{0.60} - N^{0.33}} + \alpha_3 \tag{3}$$

and calculated the ensemble average ($\langle R_h \rangle$) from[4] :

$$\langle R_h \rangle = -\ln\left(\left\langle \exp\left(-R_h^{-1}\right)\right\rangle\right)^{-1} \tag{4}$$

#### Calculating paramagnetic relaxation enhancement

We used DEER-PREdict[6] (`https://github.com/KULL-Centre/DEERpredict`) to calculate the Paramagnetic Relaxation Enhancement (PRE) from the different ensembles. DEER-PREdict builds on earlier work[7–11] to place nitroxide spin-labels on a collection of structures obtained e.g. from a simulation and calculate the average PRE. The procedure of rotamer placement and evaluation of labelling sites is analogous to the work of Polyhach et al.[10] and we here focus on how we calculate PRE observables for large structural ensembles from molecular dynamics (MD) trajectories.

The transverse relaxation enhancement rate $\Gamma_2$ between the spin-label and backbone amide is calculated from:

$$\Gamma_2 = \frac{1}{15}\left(\frac{\mu_0}{4\pi}\right)^2 \gamma_I^2 g^2 \mu_B^2 s_e\left(s_e+1\right)\left\{4J(0)+3J\left(\omega_I\right)\right\} \tag{5}$$

Here, $\mu_0$ is the vacuum permeability, $\gamma_1$ the gyromagnetic ratio of the proton, $g$ is the electrons' g-factor, $\mu_B$ is the free electron magnetic moment and $s_e$ is the paramagnetic spin number, 1/2 for nitroxide probe.

The spectral density function $J(\omega)$ is described using a 'model-free' formalism[7–9] that takes into account the overall motions in the external magnetic field as well as the internal motion of the spin label:

$$J_{SBMF}(\omega) = \left\langle r^{-6}\right\rangle\left\{\frac{S^2\tau_c}{1+\omega^2\tau_c^2} + \frac{\left(1-S^2\right)\tau_t}{1+\omega^2\tau_t^2}\right\} \tag{6}$$

The *radial* and *angular* contributions of the order parameter are separated and $S^2$ becomes:

$$S^2 \approx S^2_{angular} S^2_{radial} \tag{7}$$

with

$$S^2_{radial} = \left\langle r^{-6} \right\rangle^{-1} \left\langle r^{-3} \right\rangle^{2} \tag{8}$$

We approximate of the angular contribution of the order parameter as proposed by[7] :

$$S^2_{angular} = \frac{1}{n_p^2} \sum_{i,j}^{N} \sum_{s,t}^{n_p} \left\{ \frac{3}{2} \left( \frac{\vec{r}_{is} \cdot \vec{r}_{jt}}{r_{is} r_{jt}} \right)^2 - \frac{1}{2} \right\} p_i p_j \tag{9}$$

For a given conformation the distance between the probe nitroxide and the nucleus is calculated as,

$$\langle r^x \rangle = \frac{1}{n_p} \sum_{i}^{N} \sum_{s}^{n_p} r^x_{jis} \times p_i \tag{10}$$

$n_p$ represents the number of equivalent nuclei in the protein paramagnetic centre (here $n_p = 1$).

We calculate the relaxation enhancement rates for each frame and obtain the ensemble average by summing all rates per frame, weighted by the appropriate weight (either uniform for Boltzmann weighted ensembles or using the weights from the BME reweighting):

$$\langle \Gamma_2 \rangle = \sum_{l}^{M} \Gamma_{2,l} \times w_l \tag{11}$$

The experimental data come in the form of intensity ratios between the paramagnetic and diamagnetic sample, and we calculate this from $\Gamma_2$ using[12] :

$$\frac{I_{para}}{I_{dia}} = \frac{R_2 \exp\left(-\Gamma_2 t_d\right)}{R_2 + \Gamma_2} \tag{12}$$

where $t_d$ is the total INEPT time of the HSQC measurement.

In order to calculate PREs one has to know or estimate the reorientation (‘tumbling’) time $\tau_c$ and $\tau_t$ for the 'internal' motion of the protein. We estimated an optimal $\tau_c$ by varying its value and calculating the RMSD between experimental and calculated intensity ratios for the five probe positions:

$$RMSD = \sqrt{\frac{1}{n}\sum_{i=1}^{n}(I_i - I_{exp})^2} \tag{13}$$

where, $n$ is the number of residues with PRE data for a given spin-label. This was done for each ensemble and we obtained an optimal $\tau_c$ in the range $\approx 1-7$ns and $\tau_t = 1$ps.

To evaluate the over all improvement after refinement of the ensembles we calculated RMSD using the optimal $\tau_c$ value across the five probe-position:

$$RMSD = \sqrt{\frac{1}{n}\frac{1}{m}\sum_{i=1}^{m}\sum_{i=1}^{n}(I_i - I_{EXP})^2} \tag{14}$$

Here, $m$ is the number probe-positions ($m = 5$).

### Collective variables used in the metadynamics simulations

#### The radius of gyration ($R_{\mathrm{g}}$)

In Plumed[13] the radius of gyration ($R_{\mathrm{g}}$) is defined as:

$$R_{\mathrm{g}} = \left(\frac{\sum_i^n m_i \left|r_i - r_{\mathrm{cm}}\right|^2}{\sum_i^n m_i}\right)^{1/2} \tag{15}$$

where the positional centre of mass is defined as:

$$r_{\mathrm{cm}} = \frac{\sum_i^n r_i m_i}{\sum_i^n m_i} \tag{16}$$

The $C_\alpha$ atoms were used for the calculations.

#### Helical content (ALPHA RMSD)

These collective variables probe the $\alpha$-helical content of a protein structure, by counting how many fragments of 6 contiguous residues that form an alpha helix, and computing their RMSD distances of the backbone $N, O, C, C_\alpha$, and $C_\beta$ atoms with respect to an ideal helical conformation. The ALPHARMSD function sums up the RMSD of the distances accordingly:

$$s = \sum_i \frac{1 - \left(\frac{r_i - d_0}{r_0}\right)^n}{1 - \left(\frac{r_i - d_0}{r_0}\right)^m} \tag{17}$$

### Experiments

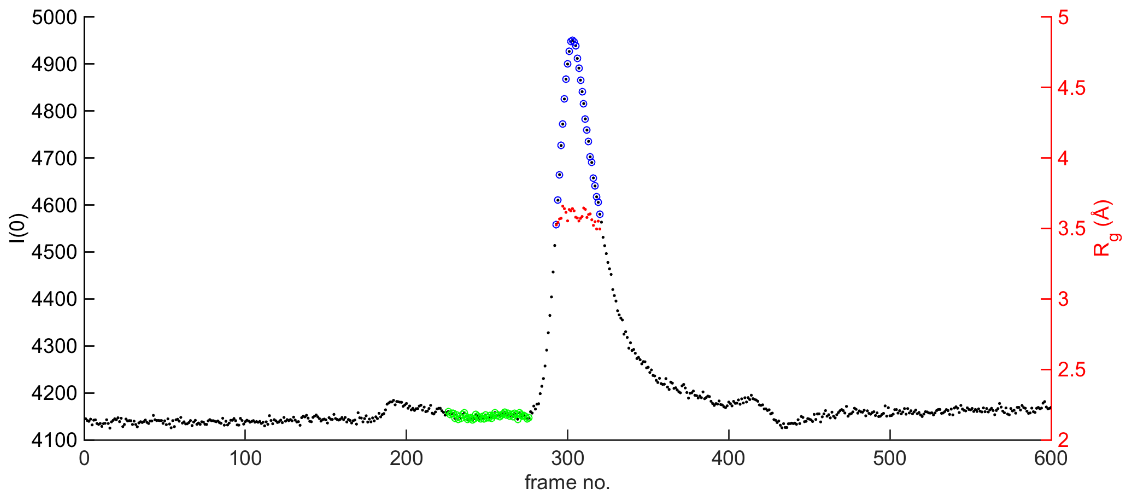


Figure 1: **SEC-SAXS experiment**. SEC-SAXS elution profile shown as 1 s frames vs. corresponding forward scattering (I(0)). Highlighted regions show measurements used for monomeric $\alpha SN$ (blue) and buffer (green), respectively. The estimated $R_g$ (red) is shown for the individual frames across the protein peak.

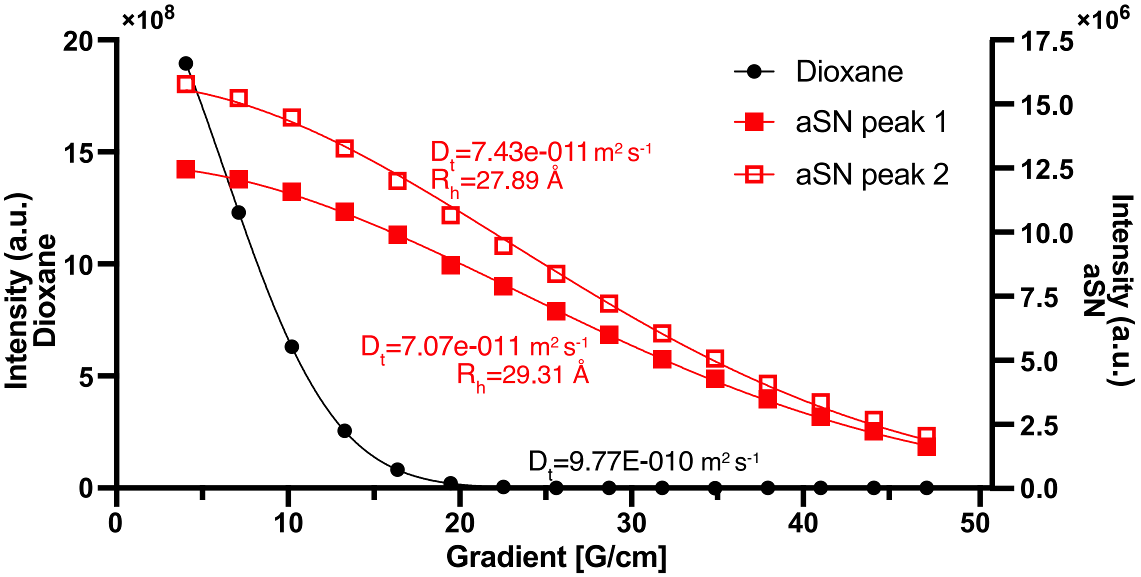


Figure 2: **NMR diffusion experiments**. NMR diffusion experiments were used to measure the (relative) translationational diffusion coefficient $D_t$ of $\alpha SN$ and dioxane as a reference. These values were used to estimate $R_h$ as described in the main text. We analysed two peaks in the NMR spectrum, which together were used to estimate $R_h = 28.6\text{Å} \pm 0.7\text{Å}$

#### Metainference results

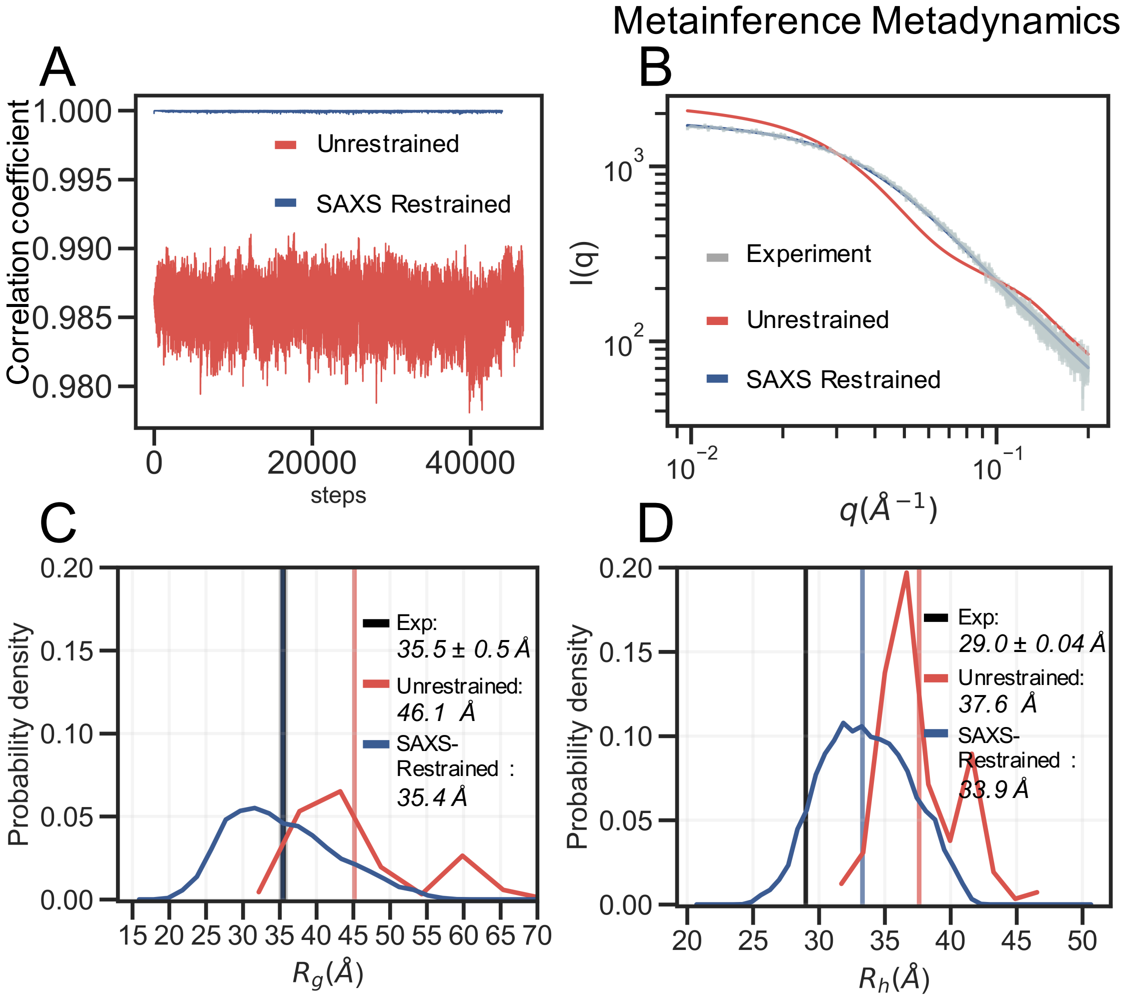


Figure 3: **Metainference Metadynamics simulation**. SAXS-restrained metainference and an unrestrained references simulation were performed with the CHARMM36m force field and the EEF1-SB implicit water model. (A) Correlation coefficient between experimental and calculated SAXS data from the (red) unrestrained and the (blue) restrained metainference simulation. (B) The calculated SAXS intensity from the (red) unrestrained and the (blue) restrained metainference simulation are compared to the (gray) experimental SAXS curve. Probability distribution of (C) $R_g$ and (D) $R_h$ for the (red) prior and (blue) reweighted ensemble. Average $R_g$ and $R_h$ and the experimental values are shown as verticals lines.

#### BME refinement

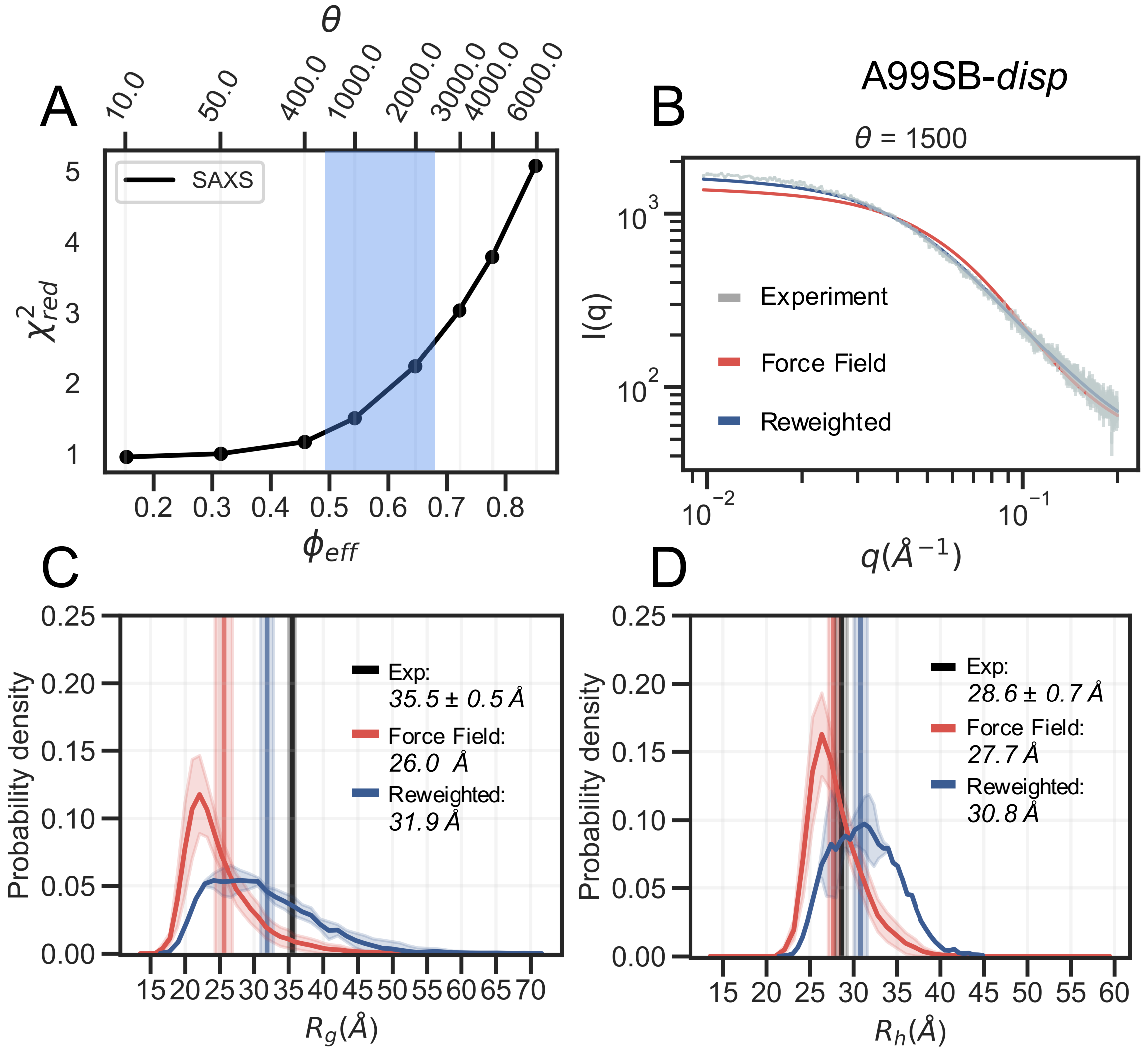


Figure 4: **BME refinement of simulation with A99SB-*disp* and with modified a TIP4P-D water model**. SAXS refinement of prior ensemble sampled with A99SB-*disp* and with modified a TIP4P-D water model. (A) L-curve selection by calculating $\chi^2_{red}$ at different values of $\theta$ and plotted against the effective fraction of frames used in reweighting ($\phi_{eff}$). For this we selected an value of $\theta = 1500$. (B) The calculated SAXS intensity from the (red) unrestrained and the (blue) restrained metainference simulation are compared to the (gray) experimental SAXS curve. Probability distribution of (C) $R_g$ and (D) $R_h$ for the (red) prior and (blue) reweighted ensemble. Average $R_g$ and $R_h$ and the experimental values are shown as verticals lines. The error of the distributions and on the averages(shown as shades) were estimated by block averaging.

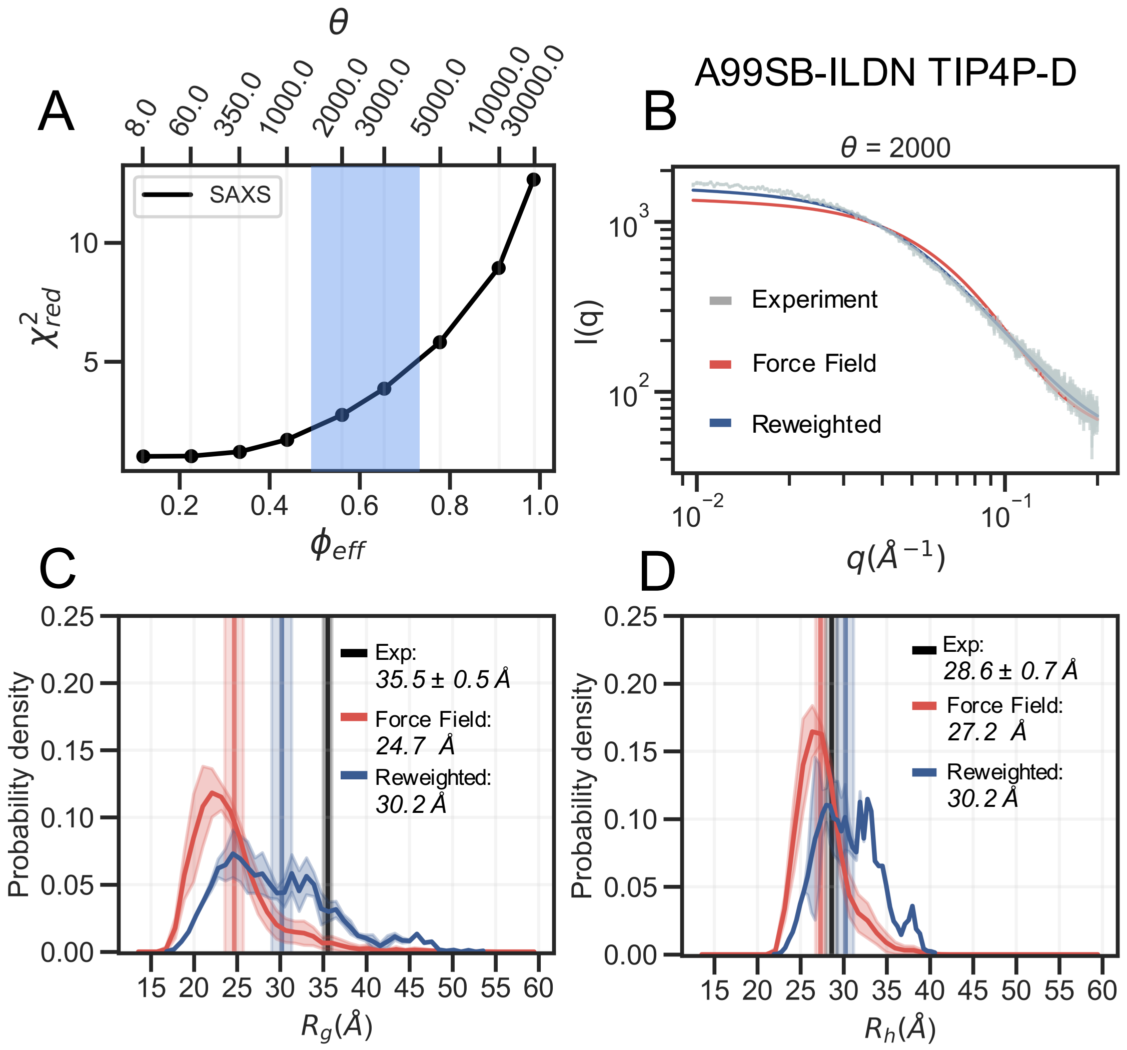


Figure 5: **BME refinement of simulation with A99SB-ILDN and with a modified TIP4P-D water model**. SAXS refinement of prior ensemble sampled with A99SB-ILDN and with a modified TIP4P-D water model. (A) L-curve selection by calculating $\chi^2_{red}$ at different values of $\theta$ and plotted against the effective fraction of frames used in reweighting ($\phi_{eff}$). For this we selected an value of $\theta = 2000$. (B) The calculated SAXS intensity from the (red) unrestrained and the (blue) restrained metainference simulation are compared to the (gray) experimental SAXS curve. Probability distribution of (C) $R_g$ and (D) $R_h$ for the (red) prior and (blue) reweighted ensemble. Average $R_g$ and $R_h$ and the experimental values are shown as verticals lines. The error of the distributions and on the averages(shown as shades) were estimated by block averaging.

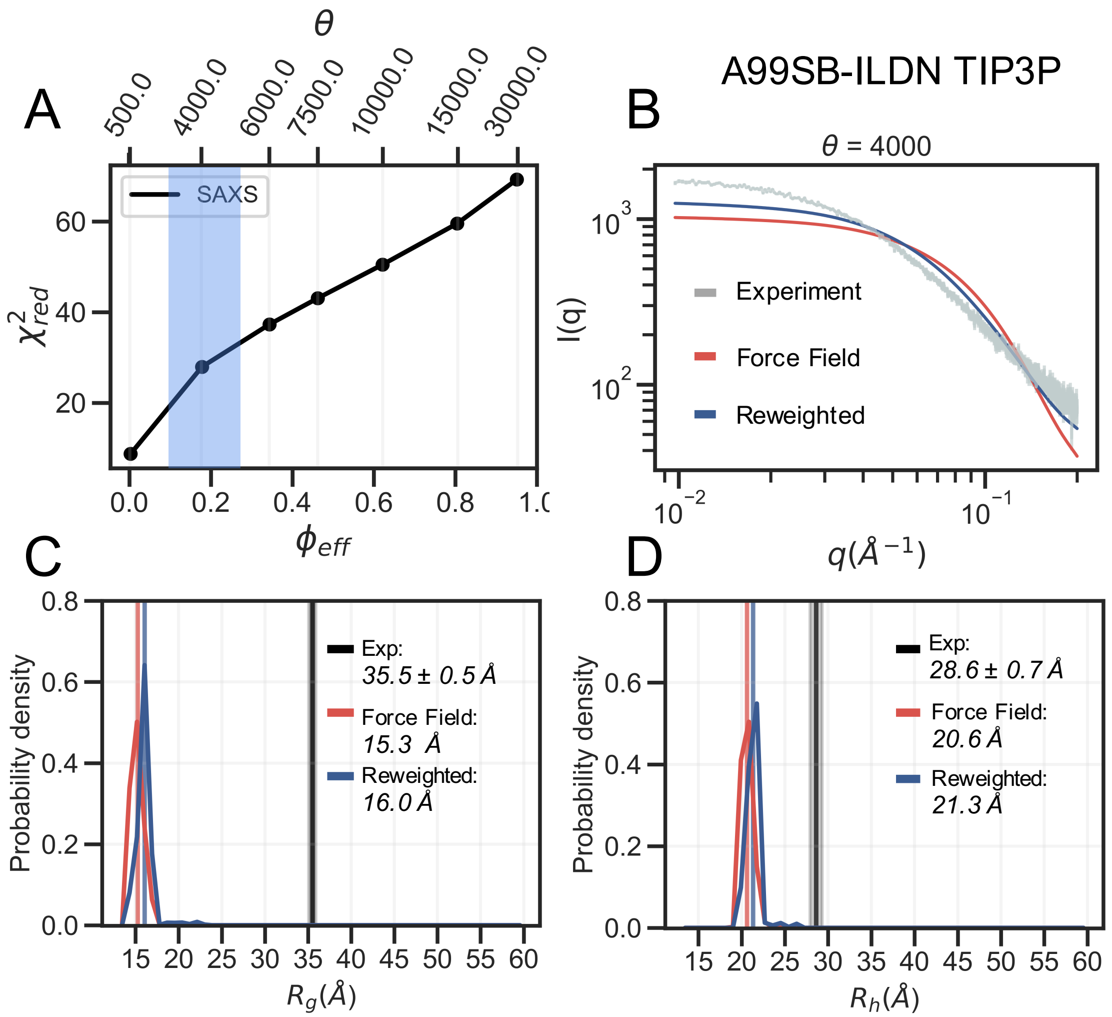


Figure 6: **BME refinement of simulation with A99SB-ILDN and with the TIP3P water model**. SAXS refinement of prior ensemble sampled with A99SB-ILDN and with the TIP3P water model. (A) L-curve selection by calculating $\chi^2_{red}$ at different values of $\theta$ and plotted against the effective fraction of frames used in reweighting ($\phi_{eff}$). For this we selected an value of $\theta = 4000$. (B) The calculated SAXS intensity from the (red) unrestrained and the (blue) restrained metainference simulation are compared to the (gray) experimental SAXS curve. Probability distribution of (C) $R_g$ and (D) $R_h$ for the (red) prior and (blue) reweighted ensemble. Average $R_g$ and $R_h$ and the experimental values are shown as verticals lines. The error of the distributions and on the averages(shown as shades) were estimated by block averaging.

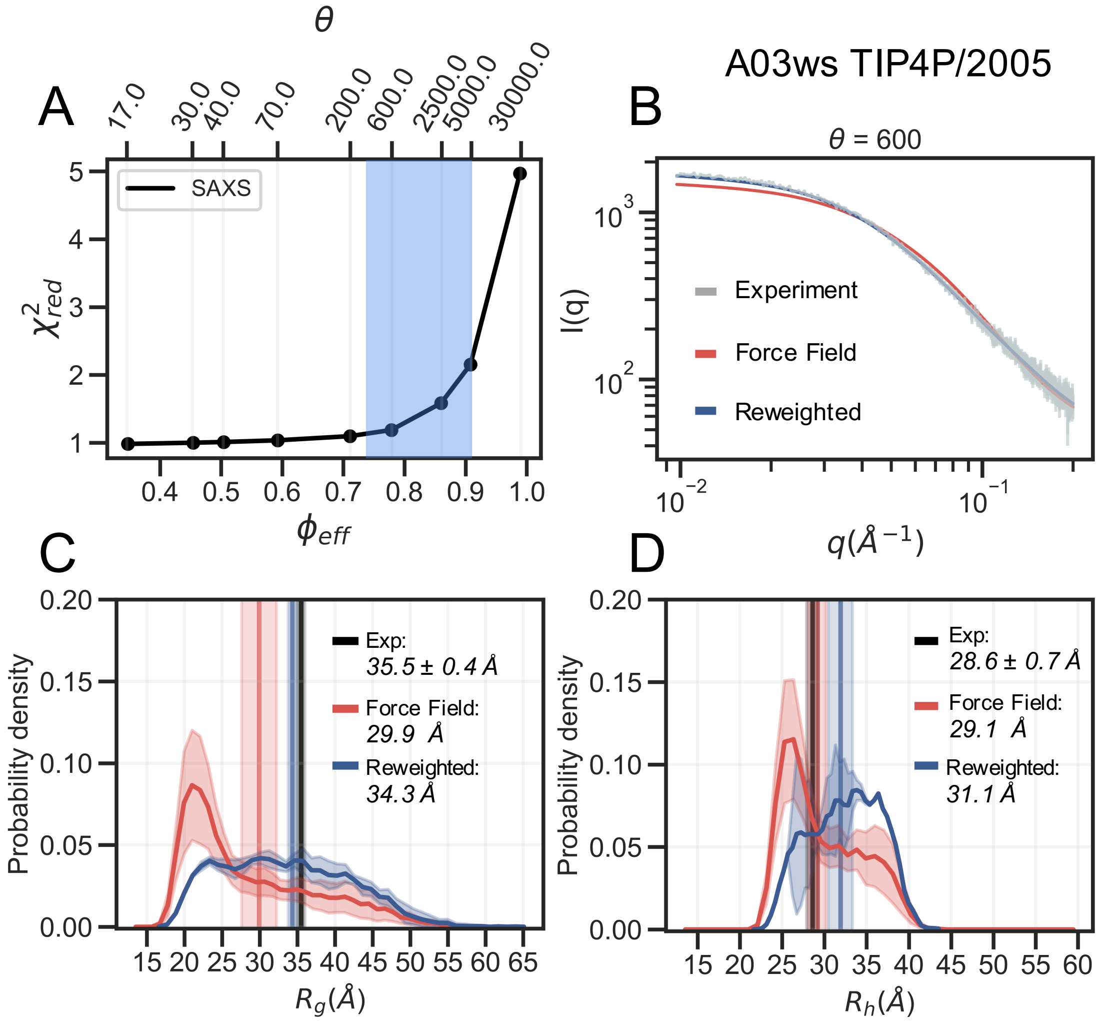


Figure 7: **BME refinement of simulation with A03ws and with the TIP4P/2005 water model**. SAXS refinement of prior ensemble sampled with A03ws and with the TIP4P/2005 water model. (A) L-curve selection by calculating $\chi^2_{red}$ at different values of $\theta$ and plotted against the effective fraction of frames used in reweighting ($\phi_{eff}$). For this we selected an value of $\theta = 600$. (B) The calculated SAXS intensity from the (red) unrestrained and the (blue) restrained metainference simulation are compared to the (gray) experimental SAXS curve. Probability distribution of (C) $R_g$ and (D) $R_h$ for the (red) prior and (blue) reweighted ensemble. Average $R_g$ and $R_h$ and the experimental values are shown as verticals lines. The error of the distributions and on the averages(shown as shades) were estimated by block averaging.

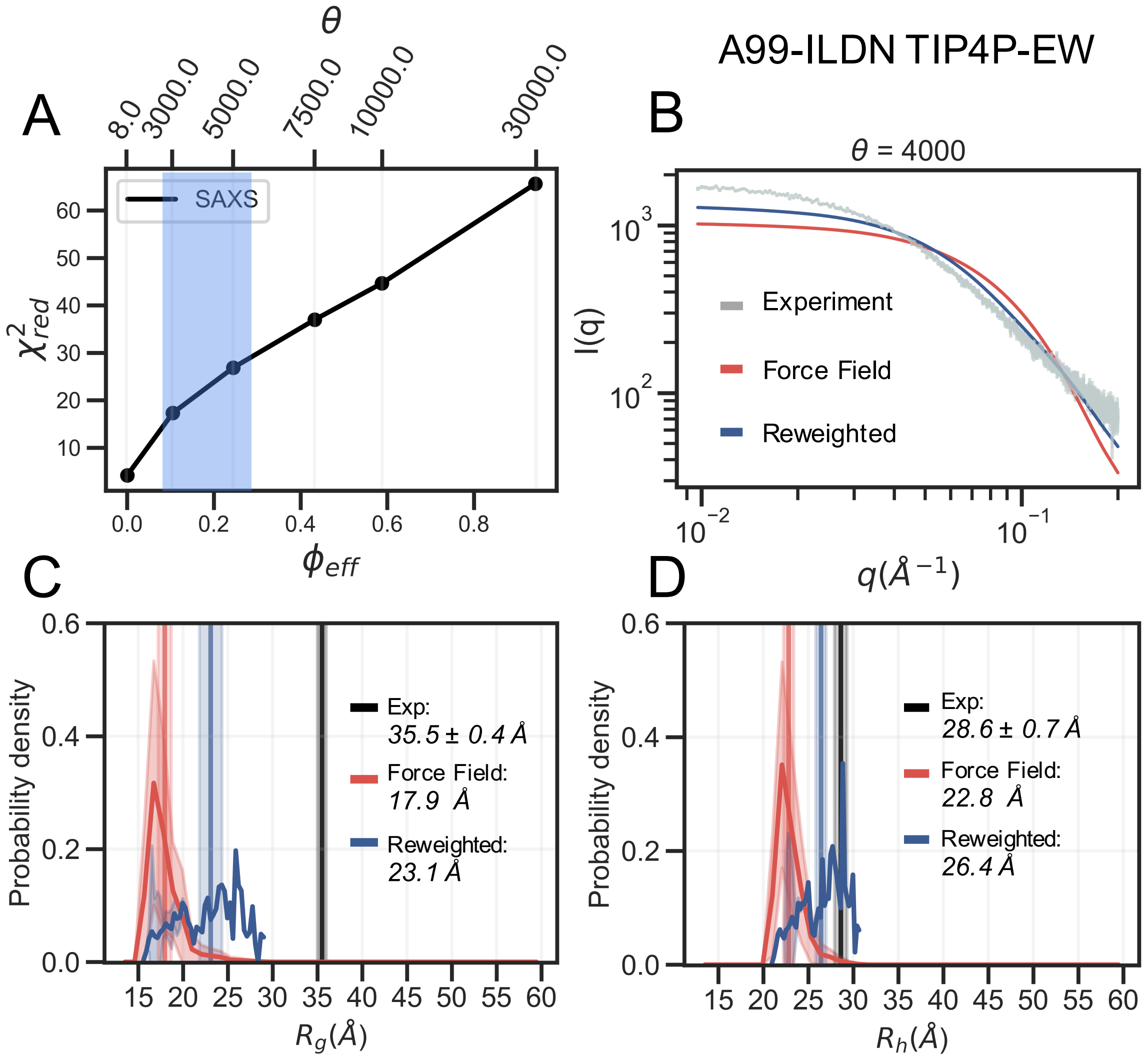


Figure 8: **BME refinement of simulation with A99-ILDN and with the TIP4P-EW water model**. SAXS refinement of prior ensemble sampled with A99-ILDN and with the TIP4P-EW water model. (A) L-curve selection by calculating $\chi^2_{red}$ at different values of $\theta$ and plotted against the effective fraction of frames used in reweighting ($\phi_{eff}$). For this we selected an value of $\theta = 4000$. (B) The calculated SAXS intensity from the (red) unrestrained and the (blue) restrained metainference simulation are compared to the (gray) experimental SAXS curve. Probability distribution of (C) $R_g$ and (D) $R_h$ for the (red) prior and (blue) reweighted ensemble. Average $R_g$ and $R_h$ and the experimental values are shown as verticals lines. The error of the distributions and on the averages(shown as shades) were estimated by block averaging.

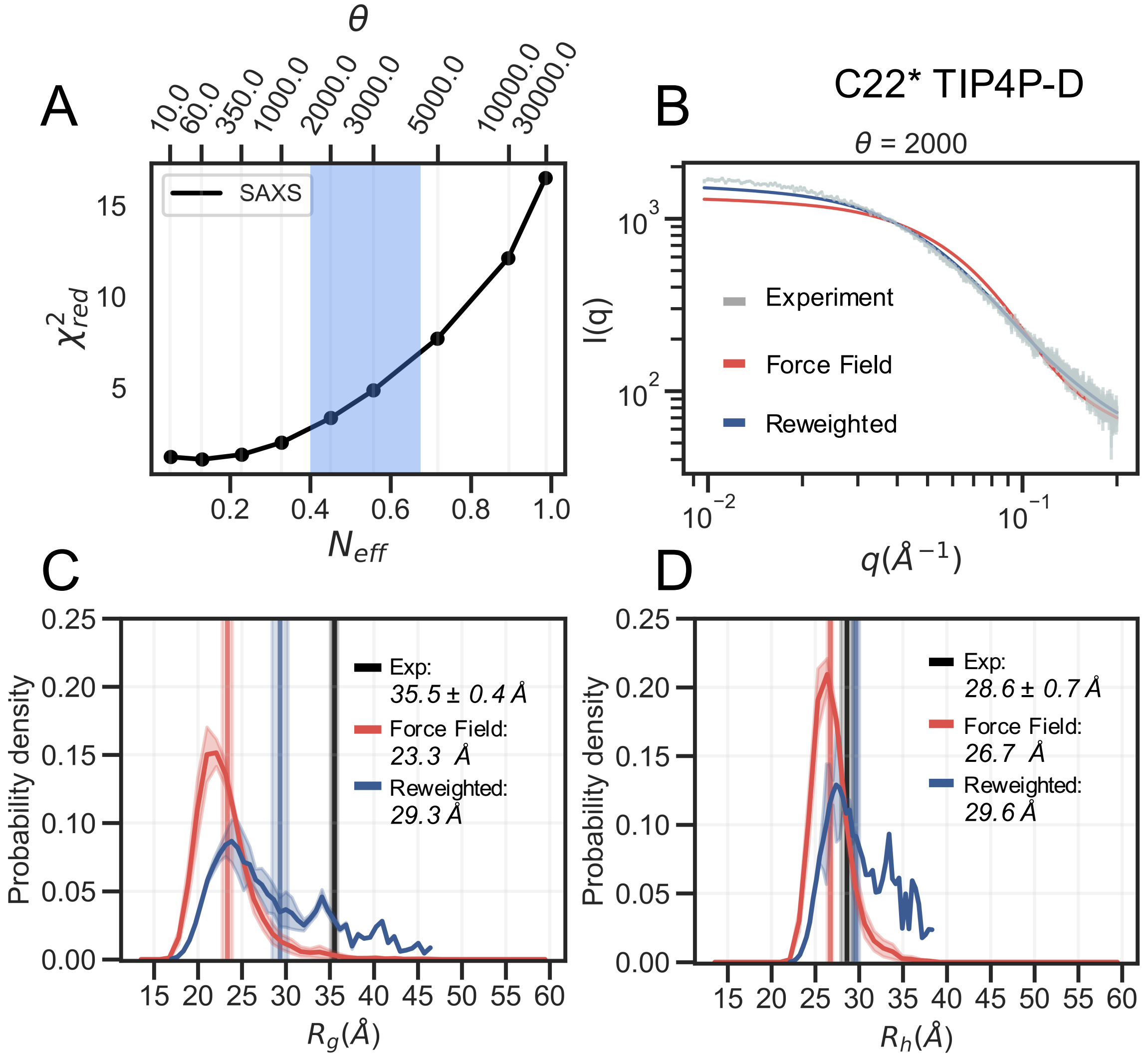


Figure 9: **BME refinement of simulation with C22* and with a modified TIP4P-D water model**. SAXS refinement of prior ensemble sampled with C22* and with a modified TIP4P-D water model. (A) L-curve selection by calculating $\chi^2_{red}$ at different values of $\theta$ and plotted against the effective fraction of frames used in reweighting ($\phi_{eff}$). For this we selected an value of $\theta = 2000$. (B) The calculated SAXS intensity from the (red) unrestrained and the (blue) restrained metainference simulation are compared to the (gray) experimental SAXS curve. Probability distribution of (C) $R_g$ and (D) $R_h$ for the (red) prior and (blue) reweighted ensemble. Average $R_g$ and $R_h$ and the experimental values are shown as verticals lines. The error of the distributions and on the averages(shown as shades) were estimated by block averaging.

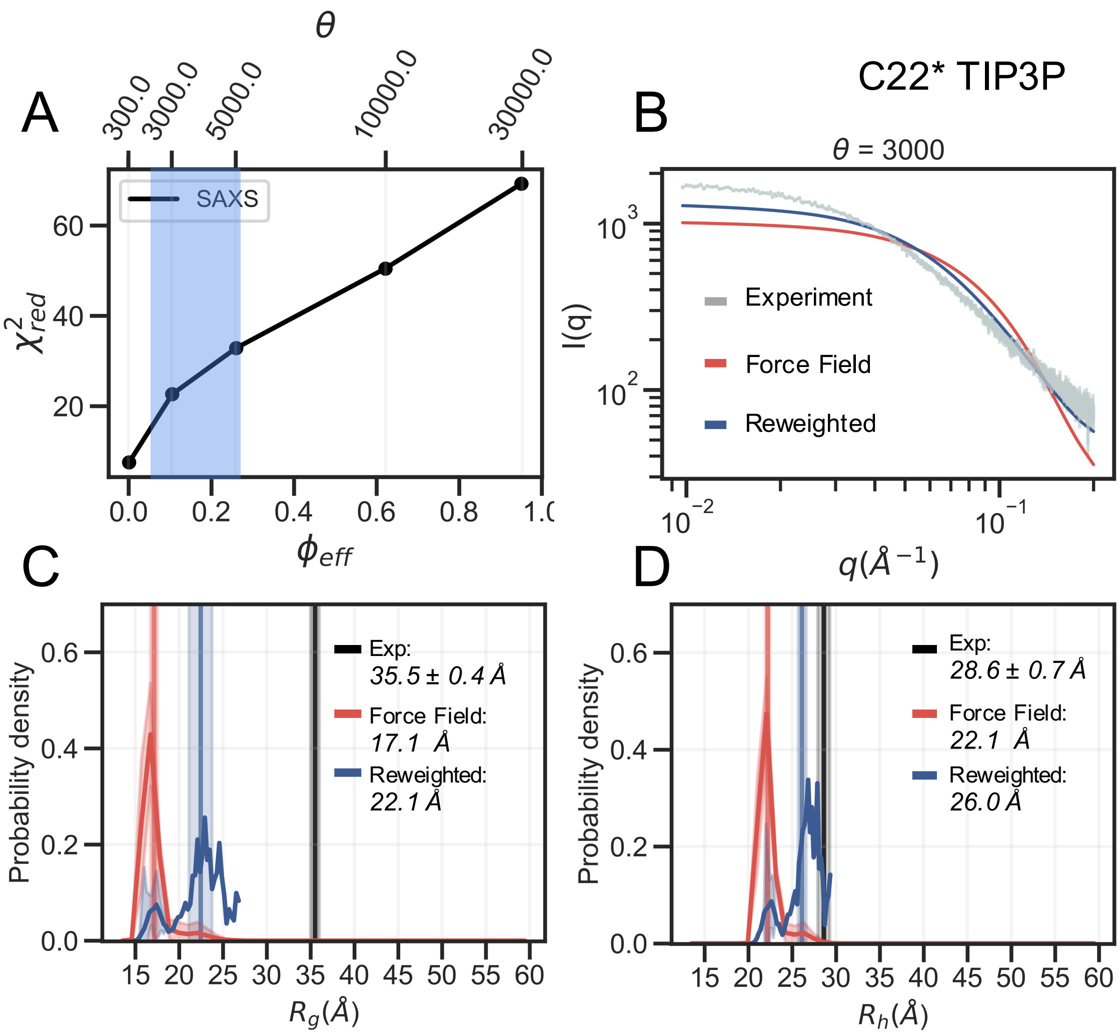


Figure 10: **BME refinement of simulation with C22* and with the TIP3P water model**. SAXS refinement of prior ensemble sampled with C22* and with the TIP3P water model. (A) L-curve selection by calculating $\chi^2_{red}$ at different values of $\theta$ and plotted against the effective fraction of frames used in reweighting ($\phi_{eff}$). For this we selected an value of $\theta = 3000$. (B) The calculated SAXS intensity from the (red) unrestrained and the (blue) restrained metainference simulation are compared to the (gray) experimental SAXS curve. Probability distribution of (C) $R_g$ and (D) $R_h$ for the (red) prior and (blue) reweighted ensemble. Average $R_g$ and $R_h$ and the experimental values are shown as verticals lines. The error of the distributions and on the averages(shown as shades) were estimated by block averaging.

#### Calculated and experimental PREs

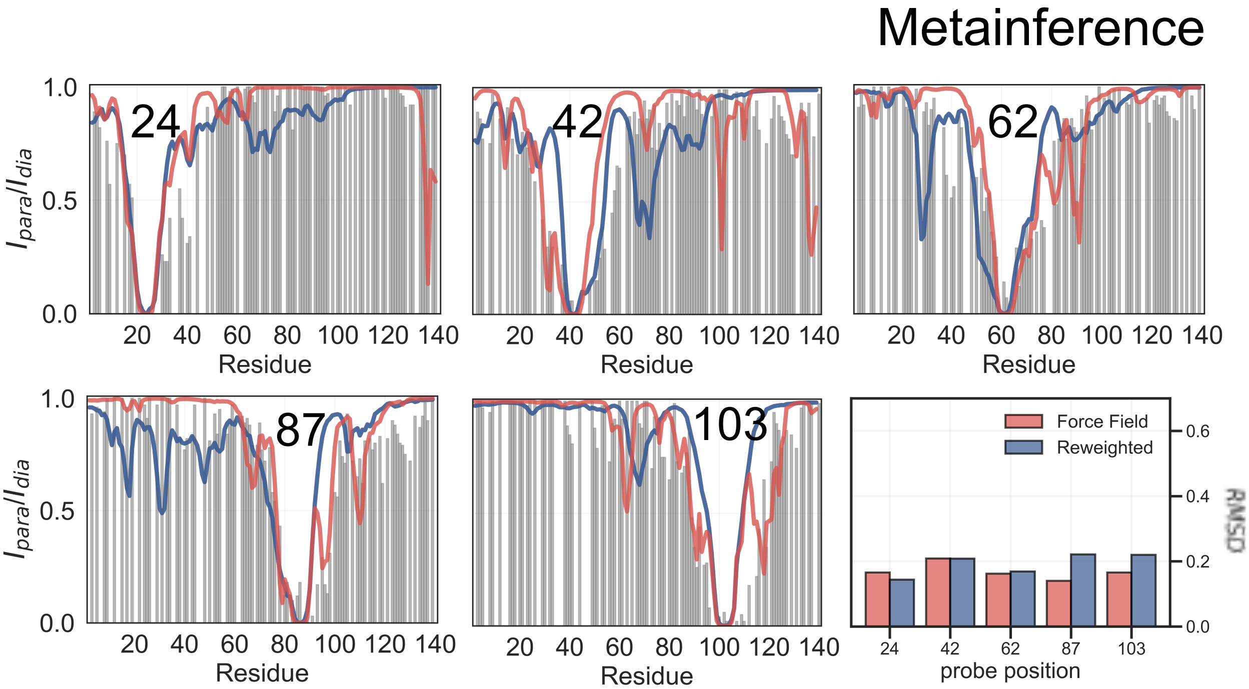


Figure 11: **Comparison of experimental PRE data with data calculated from metainference simulations with CHARMM36m force field and the EEF1-SB implicit water mode**. (Black) Experimental intensity ratios for the five probe-positions: 24,42,62,87 and 103 are compared to calculated values from (red) unrestrained and (blue) the SAXS-restrained metainference ensemble. The labelling position is denoted in each plot. The panel at the bottom right shows the RMSD (for each probe position) between the experimental values and those calculated from the (red) unrestrained and (blue) the SAXS-restrained metainference ensemble.

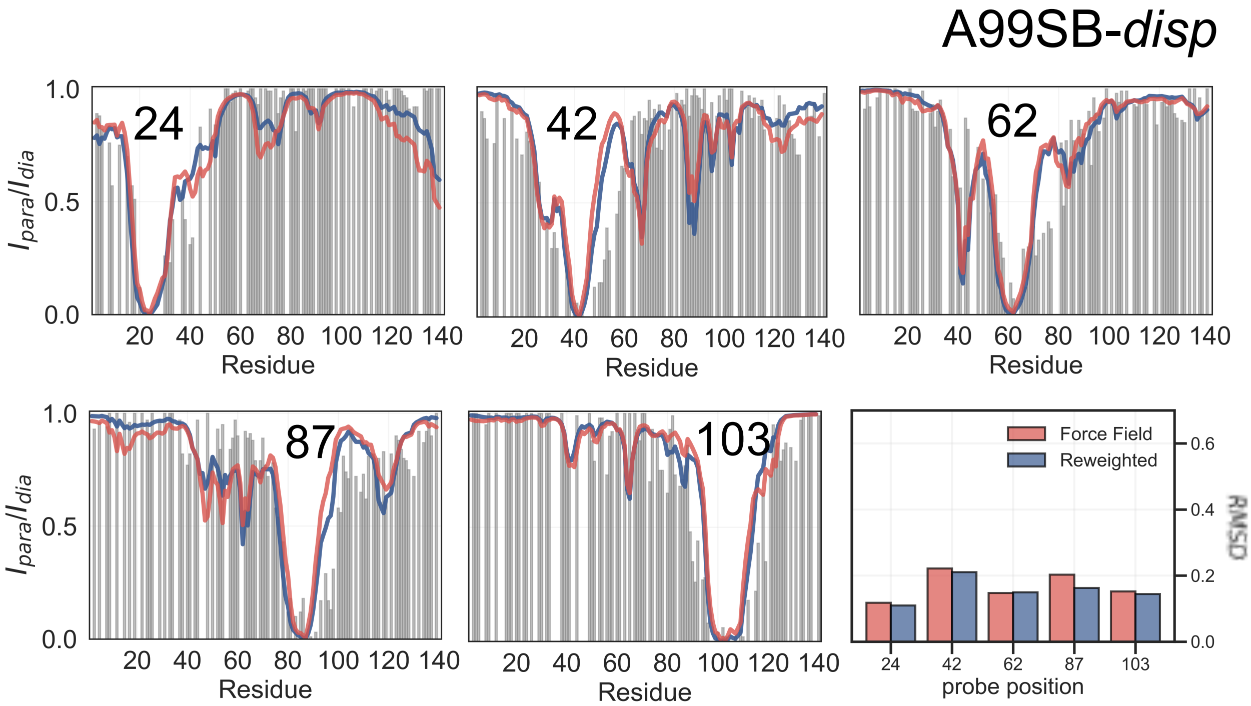


Figure 12: **Comparison of experimental PRE data with data calculated from simulations with A99SB-*disp* and with modified a TIP4P-D water model**. (Black) Experimental intensity ratios for the five probe-positions: 24,42,62,87 and 103 are compared to calculated values from MD simulations (red) before and (blue) after reweighting with SAXS data. The labelling position is denoted in each plot. The panel at the bottom right shows the RMSD (for each probe position) between the experimental values and those calculated from (red) before and (blue) after reweighting.

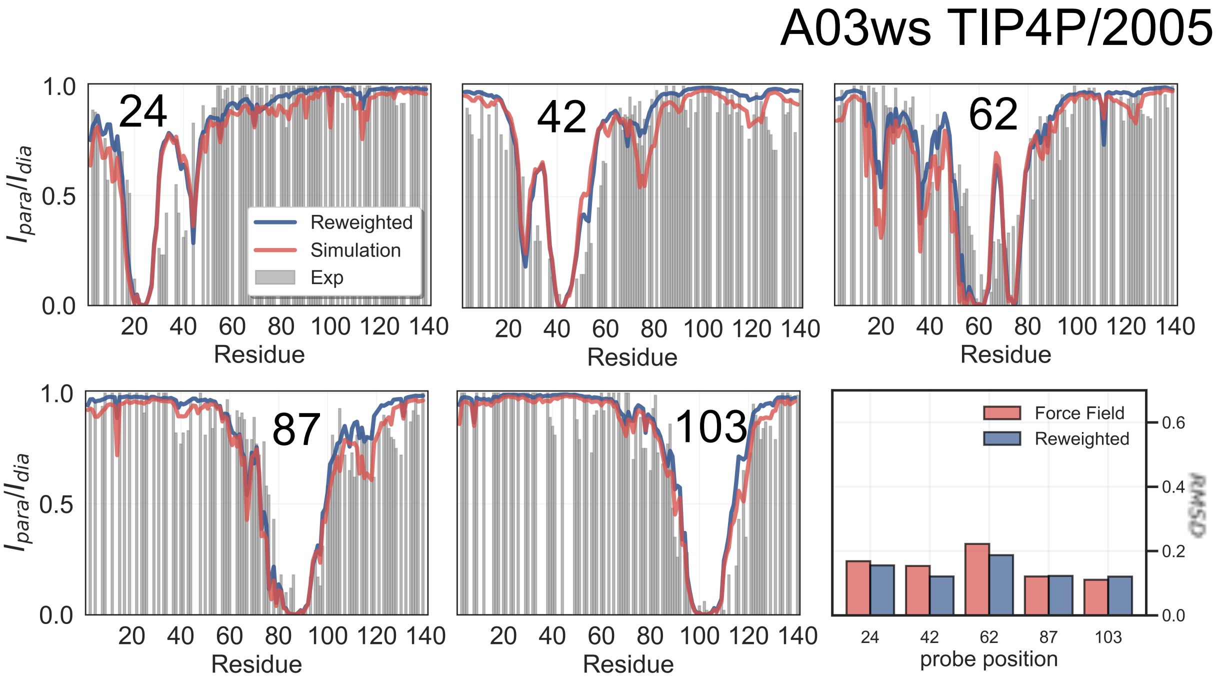


Figure 13: **Comparison of experimental PRE data with data calculated from simulations with A03ws with the TIP4P/2005 water model**. (Black) Experimental intensity ratios for the five probe-positions: 24,42,62,87 and 103 are compared to calculated values from MD simulations (red) before and (blue) after reweighting with SAXS data. The labelling position is denoted in each plot. The panel at the bottom right shows the RMSD (for each probe position) between the experimental values and those calculated from (red) before and (blue) after reweighting.

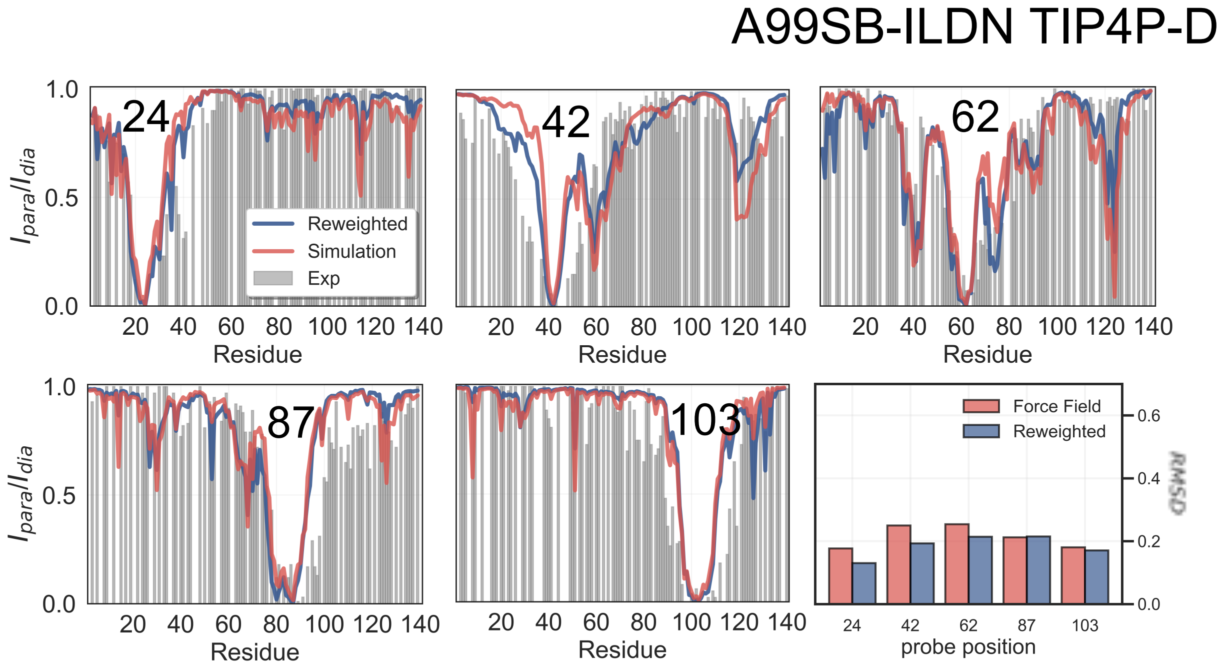


Figure 14: **Comparison of experimental PRE data with data calculated from simulations with A99SB-ILDN with the TIP4P-D water model**. (Black) Experimental intensity ratios for the five probe-positions: 24,42,62,87 and 103 are compared to calculated values from MD simulations (red) before and (blue) after reweighting with SAXS data. The labelling position is denoted in each plot. The panel at the bottom right shows the RMSD (for each probe position) between the experimental values and those calculated from (red) before and (blue) after reweighting.

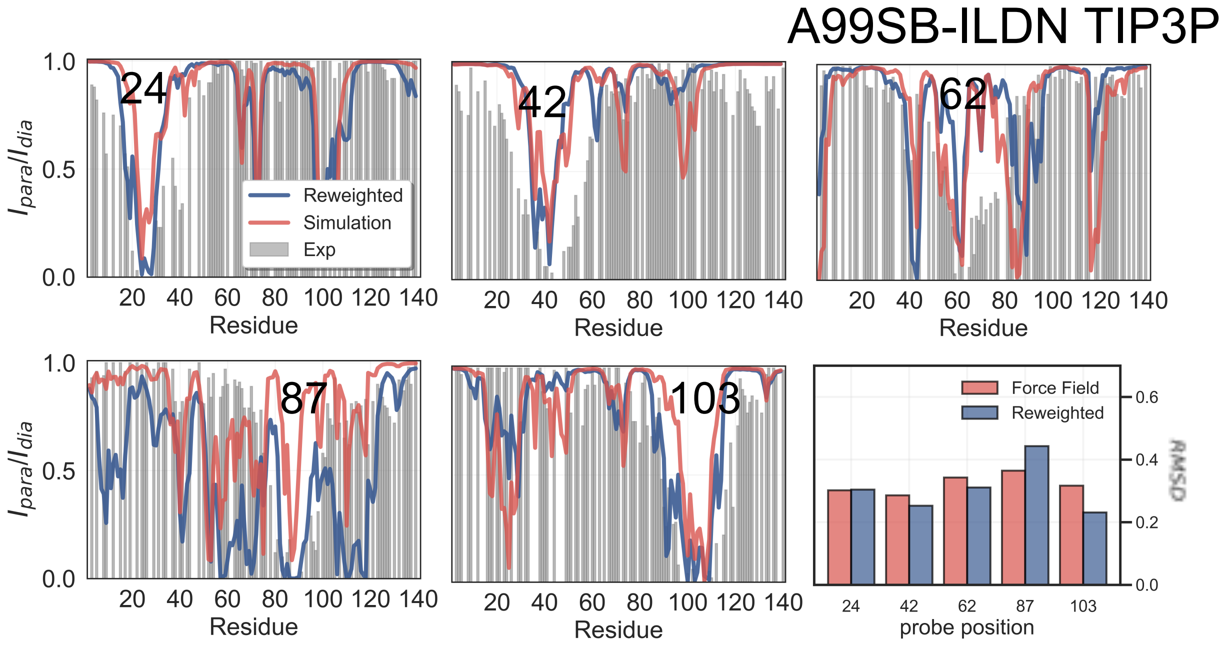


Figure 15: **Comparison of experimental PRE data with data calculated from simulations with A99SB-ILDN with the TIP3P water model**. (Black) Experimental intensity ratios for the five probe-positions: 24,42,62,87 and 103 are compared to calculated values from MD simulations (red) before and (blue) after reweighting with SAXS data. The labelling position is denoted in each plot. The panel at the bottom right shows the RMSD (for each probe position) between the experimental values and those calculated from (red) before and (blue) after reweighting.

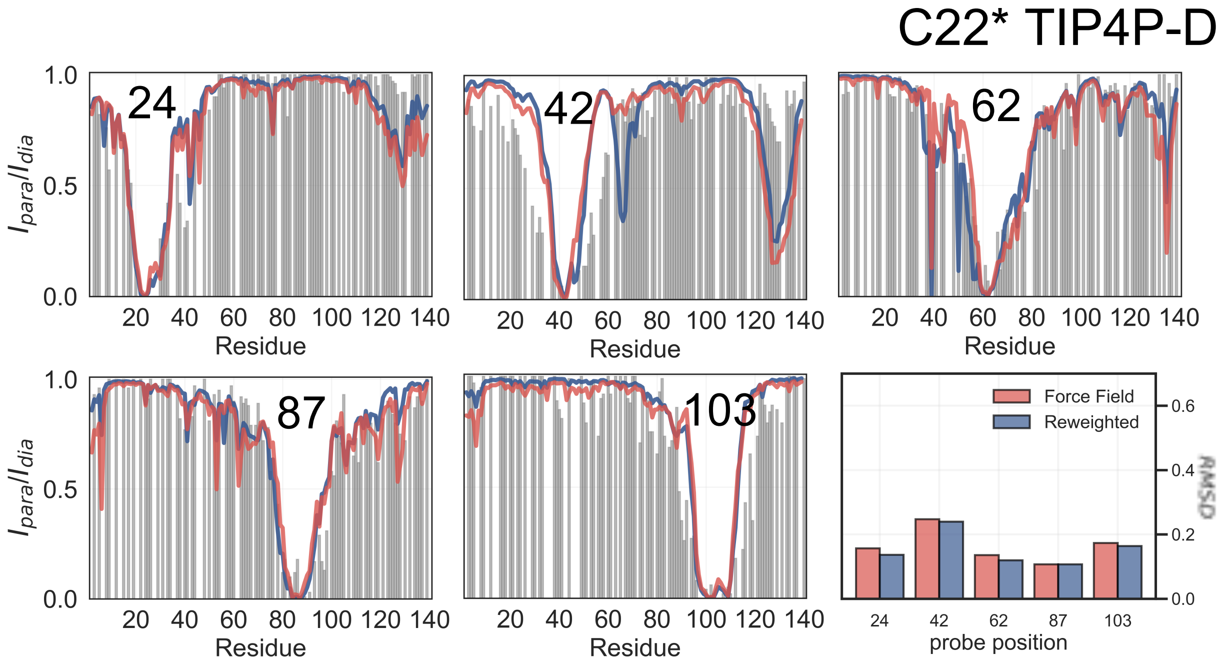


Figure 16: **Comparison of experimental PRE data with data calculated from simulations with C22* with the TIP4P-D water model**. (Black) Experimental intensity ratios for the five probe-positions: 24,42,62,87 and 103 are compared to calculated values from MD simulations (red) before and (blue) after reweighting with SAXS data. The labelling position is denoted in each plot. The panel at the bottom right shows the RMSD (for each probe position) between the experimental values and those calculated from (red) before and (blue) after reweighting.

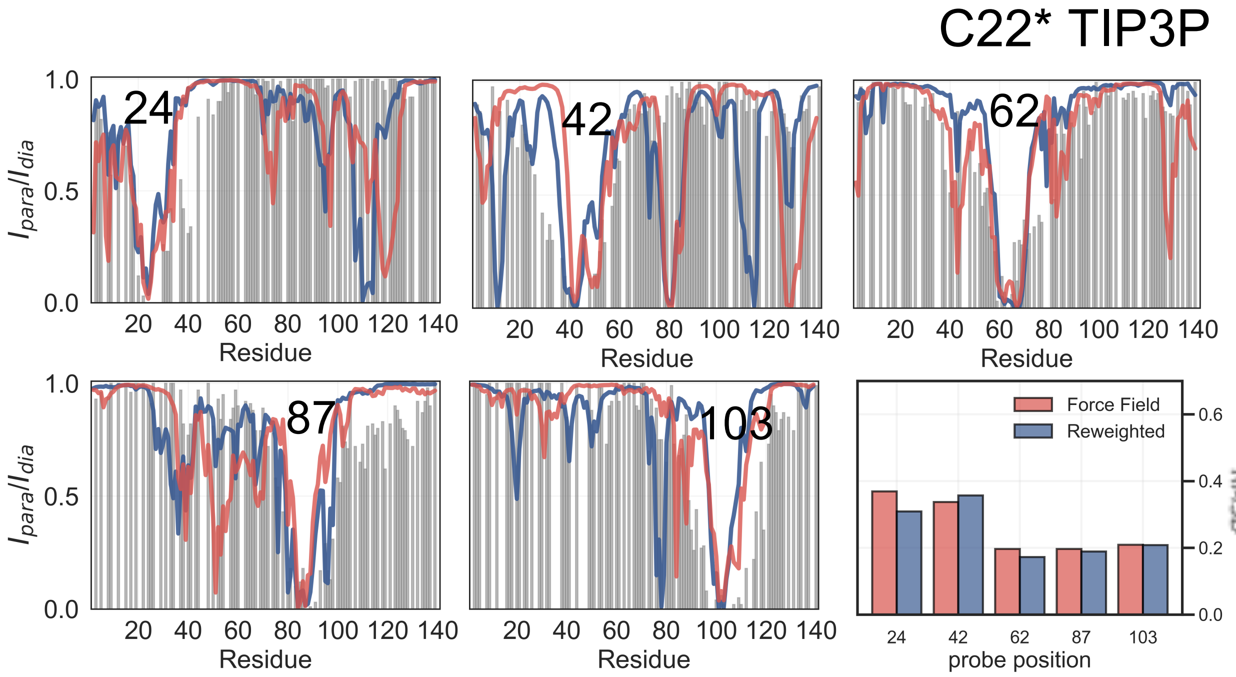


Figure 17: **Comparison of experimental PRE data with data calculated from simulations with C22* with the TIP3P water model**. (Black) Experimental intensity ratios for the five probe-positions: 24,42,62,87 and 103 are compared to calculated values from MD simulations (red) before and (blue) after reweighting with SAXS data. The labelling position is denoted in each plot. The panel at the bottom right shows the RMSD (for each probe position) between the experimental values and those calculated from (red) before and (blue) after reweighting.

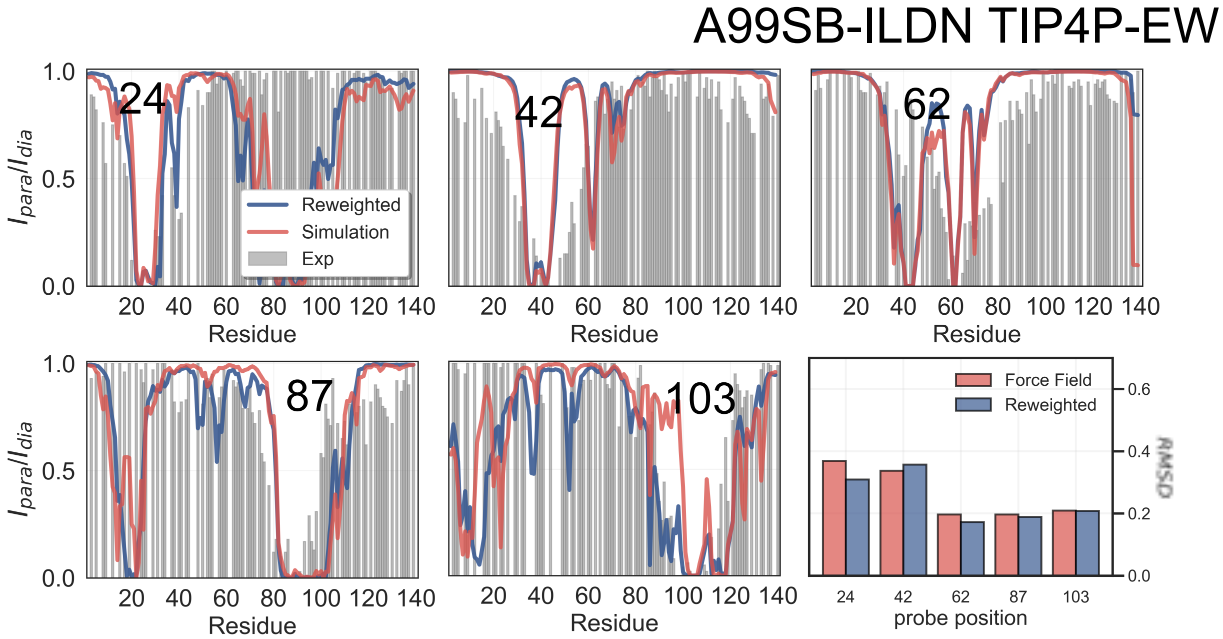


Figure 18: **Comparison of experimental PRE data with data calculated from simulations with A99SB-ILDN with TIP4P-EW water model**. (Black) Experimental intensity ratios for the five probe-positions: 24,42,62,87 and 103 are compared to calculated values from MD simulations (red) before and (blue) after reweighting with SAXS data. The labelling position is denoted in each plot. The panel at the bottom right shows the RMSD (for each probe position) between the experimental values and those calculated from (red) before and (blue) after reweighting.

### References


(1) Grudinin, S.; Garkavenko, M.; Kazennov, A. Pepsi-SAXS: an adaptive method for rapid and accurate computation of small-angle X-ray scattering profiles. *Acta Crystallographica Section D: Structural Biology* **2017**, *73*, 449–464.

(2) Bengtsen, T.; Holm, V. L.; Kjølbye, L. R.; Midtgaard, S. R.; Johansen, N. T.; Tesei, G.; Bottaro, S.; Schiøtt, B.; Arleth, L.; Lindorff-Larsen, K. Structure and dynamics of a nanodisc by integrating NMR, SAXS and SANS experiments with molecular dynamics simulations. *Elife* **2020**, *9*, e56518.

(3) Larsen, A. H.; Wang, Y.; Bottaro, S.; Grudinin, S.; Arleth, L.; Lindorff-Larsen, K. Combining molecular dynamics simulations with small-angle X-ray and neutron scattering data to study multi-domain proteins in solution. *PLoS computational biology* **2020**, *16*, e1007870.

(4) Ahmed, M. C.; Crehuet, R.; Lindorff-Larsen, K. *Intrinsically Disordered Proteins*; Springer, 2020; pp 429–445.

(5) Nygaard, M.; Kragelund, B. B.; Papaleo, E.; Lindorff-Larsen, K. An efficient method for estimating the hydrodynamic radius of disordered protein conformations. *Biophysical journal* **2017**, *113*, 550–557.

(6) Tesei, G.; Martins, J. M.; Kunze, M. B.; Wang, Y.; Crehuet, R.; Lindorff-Larsen, K. DEER-PREdict: software for efficient calculation of Spin-Labeling EPR and NMR data from conformational ensembles. *bioRxiv* **2020**,

(7) Brüschweiler, R.; Roux, B.; Blackledge, M.; Griesinger, C.; Karplus, M.; Ernst, R. Influence of rapid intramolecular motion on NMR cross-relaxation rates. A molecular dynamics study of antamanide in solution. *Journal of the American Chemical Society* **1992**, *114*, 2289–2302.

(8) Iwahara, J.; Schwieters, C. D.; Clore, G. M. Ensemble approach for NMR structure refinement against 1H paramagnetic relaxation enhancement data arising from a flexible paramagnetic group attached to a macromolecule. *Journal of the American Chemical Society* **2004**, *126*, 5879–5896.

(9) Salmon, L.; Nodet, G.; Ozenne, V.; Yin, G.; Jensen, M. R.; Zweckstetter, M.; Blackledge, M. NMR characterization of long-range order in intrinsically disordered proteins. *Journal of the American Chemical Society* **2010**, *132*, 8407–8418.

(10) Polyhach, Y.; Bordignon, E.; Jeschke, G. Rotamer libraries of spin labelled cysteines for protein studies. *Physical Chemistry Chemical Physics* **2011**, *13*, 2356–2366.

(11) Klose, D.; Klare, J. P.; Grohmann, D.; Kay, C. W.; Werner, F.; Steinhoff, H.-J. Simulation vs. reality: a comparison of in silico distance predictions with DEER and FRET measurements. *PLoS One* **2012**, *7*, e39492.

(12) Battiste, J. L.; Wagner, G. Utilization of site-directed spin labeling and high-resolution heteronuclear nuclear magnetic resonance for global fold determination of large proteins with limited nuclear overhauser effect data. *Biochemistry* **2000**, *39*, 5355–5365.

(13) Tribello, G. A.; Bonomi, M.; Branduardi, D.; Camilloni, C.; Bussi, G. PLUMED 2: New feathers for an old bird. *Computer Physics Communications* **2014**, *185*, 604–613.
